# Transcriptome profiling of the Olig2-expressing astrocyte subtype reveals their unique molecular signature

**DOI:** 10.1101/2020.10.15.340505

**Authors:** David Ohayon, Marion Aguirrebengoa, Nathalie Escalas, Cathy Soula

## Abstract

Astrocytes are recognized to be a heterogeneous population of cells that differ morphologically, functionally and molecularly. Whether this heterogeneity results from generation of distinct astrocyte cell lineages, each functionally specialized to perform specific tasks, remains an open question. In this study, we used RNA-seq analysis to determine the global transcriptome profile of the Olig2-expressing astrocyte subtype (Olig2-AS), a specific spinal astrocyte subtype which segregates early during development from Olig2 progenitors and differs from other spinal astrocytes by the expression of Olig2. We identified 245 differentially expressed genes. Among them, 135 exhibit higher levels of expression when compared to other populations of spinal astrocytes, indicating that these genes can serve as a ‘unique’functional signature of Olig2-AS. Further analysis highlighted, in particular, enrichment in Olig2-AS of a set of genes specialized in regulating synaptic activity. Our work thus reveals that Olig2 progenitors produce a unique astrocyte subtype specialized to perform certain specific functions.

## Introduction

Astrocytes are the largest class of glial cells in the mammalian central nervous system (CNS) and are recognized as essential for a wide variety of complex functions. Astrocytes supply energy metabolites to neurons, regulate the blood flow and the blood-brain barrier and control the levels of extracellular ions, neurotransmitters and fluids (Allen and al., 2018). The expression of functional receptors on their plasma membrane also allows astrocytes to sense neurotransmitters from nearby synaptic sites and to respond to neurons by the release of various gliotransmitters which can influence neuronal and synaptic functions (Allen and Eroglu, 2017). A major still unsolved issue regarding astrocytes is whether the diversity of functions they assume is shared by all astrocytes or is achieved by specialized astrocyte subtypes dedicated to perform specific functions in the CNS. Based on their morphology and location, astrocytes have long been divided into two distinct classes, protoplasmic astrocytes found in the grey matter and fibrous astrocytes found in the white matter. However, interest for these cells over the last decades has provided an increasing amount of evidence for a far greater heterogeneity of astrocytes (Zhang and Barres, 2010). Based on differences in morphology and marker protein expression, different populations of astrocytes, displaying both inter- and intra-regional differences, have been identified in the grey and white matters of the rodent brain and spinal cord (Chaboub and Deneen 2013; Khakh and Sofroniev, 2015; Ben Haim and Rowitch, 2017; Pestana et al., 2020). Recently, genome-wide transcriptome profiles of astrocytes provided additional insights into astrocyte heterogeneity, resulting in an unprecedented amount of data that offer opportunities to study astrocytes phenotypes and functions in health and disease (Pestana et al., 2020). These studies revealed molecular diversity of astrocytes, across brain regions (Doyle et al., 2008; Morel et al. 2017; Boisvert et al. 2018; Clarke et al. 2018; Zeisel et al., 2018; Batiuk et al 2020, Lozzi et al., 2020) but also within regions (Zeisel et al., 2015; Gokce et al., 2016; Chai et al, 2017; Lin et al., 2017; Saunders et al, 2018; Morel et al., 2019; Bayraktar et al., 2020; Batiuk et al 2020). Gene expression profiling also revealed regionally distinct molecular differences in astrocytes populating the dorsal and ventral spinal cord (Molofsky et al. 2014). However, there still remain major gaps in our understanding about the identification and overall characterization of different astroglial cell populations.

Developmental studies, by revealing that positional identity during embryogenesis is an organizing feature of astrocyte diversity, have also proven to be very insightful for our understanding of origin and possible function of astrocyte heterogeneity. Pioneer studies performed in the embryonic spinal cord identified that distinct astrocyte subtypes are generated from separate classes of progenitor cells specified early during patterning of the neural tube (Muroyama et al. 2005; Hochstim et al. 2008). During this process, morphogen factors regulate expression of sets of transcription factors throughout the dorsoventral axis of the neural tube, thus subdividing it into distinct neural progenitor domains, each dedicated to generate specific neuronal subtypes (Dessaud et al., 2008). Gliogenesis is initiated later and, at these later phases, domain organization of the spinal cord still plays an instructive role for the generation of glial cell subtypes (Ben Haim& Rowitch, 2017). This principle was first established from the observation that oligodendrocyte precursor cells (OPC) first emerge from a discrete region in the ventral neural tube, named the pMN domain, which, earlier on, gives rise to motor neurons (MN,). Progenitor cells of the pMN domain are characterized by the expression of the transcription factor Olig2 which is required for establishment of the domain but is also recognized as a cell-intrinsic determinant essential for the specification of both MNs and OPCs (Rowitch and Kriegstein, 2010). Astrocytes were first recognized to arise from pMN adjacent but non overlapping progenitor domains and three subtypes of white matter spinal astrocytes, termed VA1, VA2, and VA3, originating from separate domains of the ventral progenitor zone, have been identified (Bayraktar et al., 2015). Importantly, lineage tracing studies have shown that astrocyte precursors (APs) migrate in a restricted segmental fashion and are regionally allocated into the spinal cord according to the dorso-ventral position of their domains of origin (Tsai et al. 2012). A similar regional specification mechanism linked to the ultimate astrocyte spatial position in the adult CNS has been shown to occur in the forebrain (Bayraktar et al., 2015). However, the question remains as to whether patterning mechanisms might establish a template for generation of the functional diversity of mature astrocytes. In favour of this assumption, region restricted astrocyte subsets have been reported to assume specialized functions in the ventral spinal cord (Molofsky et al., 2014; Kelley et al., 2018).

In a previous work, we identified a regionalized astrocyte sub-type populating the grey matter of the spinal cord that we named the Olig2-AS, for Olig2-astrocytes because they originate from Olig2 progenitor cells of the pMN domain, as do MNs and OPCs, and also because they retain expression of Olig2 as they differentiate into mature astrocytes (Ohayon et al., 2019). An important aspect of the identification of Olig2-AS is that these astrocytes can be distinguished from other spinal astrocytes at post-natal stages by the expression of Olig2. Thus, their identification offered a unique opportunity to study molecular characteristics of one specific subtype of spinal cord astrocytes. In this study, we developed a fluorescence-activated cell sorting (FACS)-based approach to purify Olig2-AS from other astrocyte populations as well as from oligodendroglial cells. We then used RNA-Seq to generate high-resolution transcriptome databases and identify the molecular signature of the Olig2-AS. Overall, our study reveals new molecular insight into the nature of astrocyte diversity in the spinal cord and highlights functional specificity of Olig2-AS in regulating synaptic activity.

## Results

### Selective labeling of Olig2-AS by expression of fluorescent reporter proteins in double transgenic mice

The prerequisite for characterizing further Olig2-AS cells was the need to isolate them from the two other main glial cell populations, namely the astrocytes that do not express Olig2 (nonOlig2-AS) and cells of the oligodendroglial lineage, i.e. OPC and differentiated oligodendocytes (OL). For this purpose, we developed a double transgenic mouse model in which all the three cell types can be distinguished by specific expression of fluorescent reporter proteins. In this model, we took advantage of the combined expression of Aldh1L1 and Olig2 in the Olig2-AS to specifically color code these cells (Ohayon et al., 2019). We thus turned to the transgenic *aldhlLl*-eGFP mouse line in which *aldhlLl* promoter drives eGFP targeted expression in all astrocytes (Cahoy et al., 2008; Heintz et al., 2004) and to an *olig2-*tdTomato (tdT) mouse line that expresses the red fluorescent protein under the control of *olig2* regulatory sequences (Jackson Laboratory). Because the latter had not yet been characterized, we first compared expression of tdT with that of the endogenous Olig2 protein performing immunodetection of Olig2 on E13.5 embryonic and P7 post-natal spinal cord sections. In these experiments, we also used Sox10, an early and reliable marker of OPC and OL in mouse (Kuhlbrodt et al., 1998; Zhou et al., 2000; Stolt et al., 2002). At E13.5, Olig2 expression was detected in pMN progenitor cells as well as in two distinct cell populations that have emigrated in the mantle zone, comprising Sox10+ OPC and Olig2-AS that do not express Sox10 (Figure S1A, Ohayon et al., 2019). We found that most, if not all, mantle zone tdT+ cells were positive for the Olig2 immunostaining (Figure S1A’-A”‘). Similarly, at P7, Olig2 immunostaining colocalized with tdT and cells co-expressing tdT and Olig2 but not Sox10 were detected in the grey matter (Figure S1B-C”‘).The *olig2-*tdTomato mouse thus faithfully reports endogenous expression of Olig2, validating the use of this mouse line to label all spinal cord Olig2-expressing cells, including the Olig2-AS. We then crossed these mice with *aldh1L1-*eGFP individuals and examined expression patterns of the reporter proteins. In these mice, Olig2-AS were expected to co-express tdT and eGFP while nonOlig2-AS and OPC/OL were expected to only express eGFP and tdT, respectively. At E18.5, a stage where Olig2-AS have already been produced in the ventral spinal cord (Ohayon et al., 2019), we were able to distinguish tdT+, eGFP+ and tdT+/eGFP+ cells in the ventral grey matter (Figure 1A-A”). The same three cell populations were detected at P7 (Figure 1B, C). Confirming that tdT+/eGFP+ cells indeed correspond to the Olig2-AS, we found that these cells were immunostained for Olig2 but not for Sox10 (Figure 1B”-b”, C”-c”).

**Figure 1:**
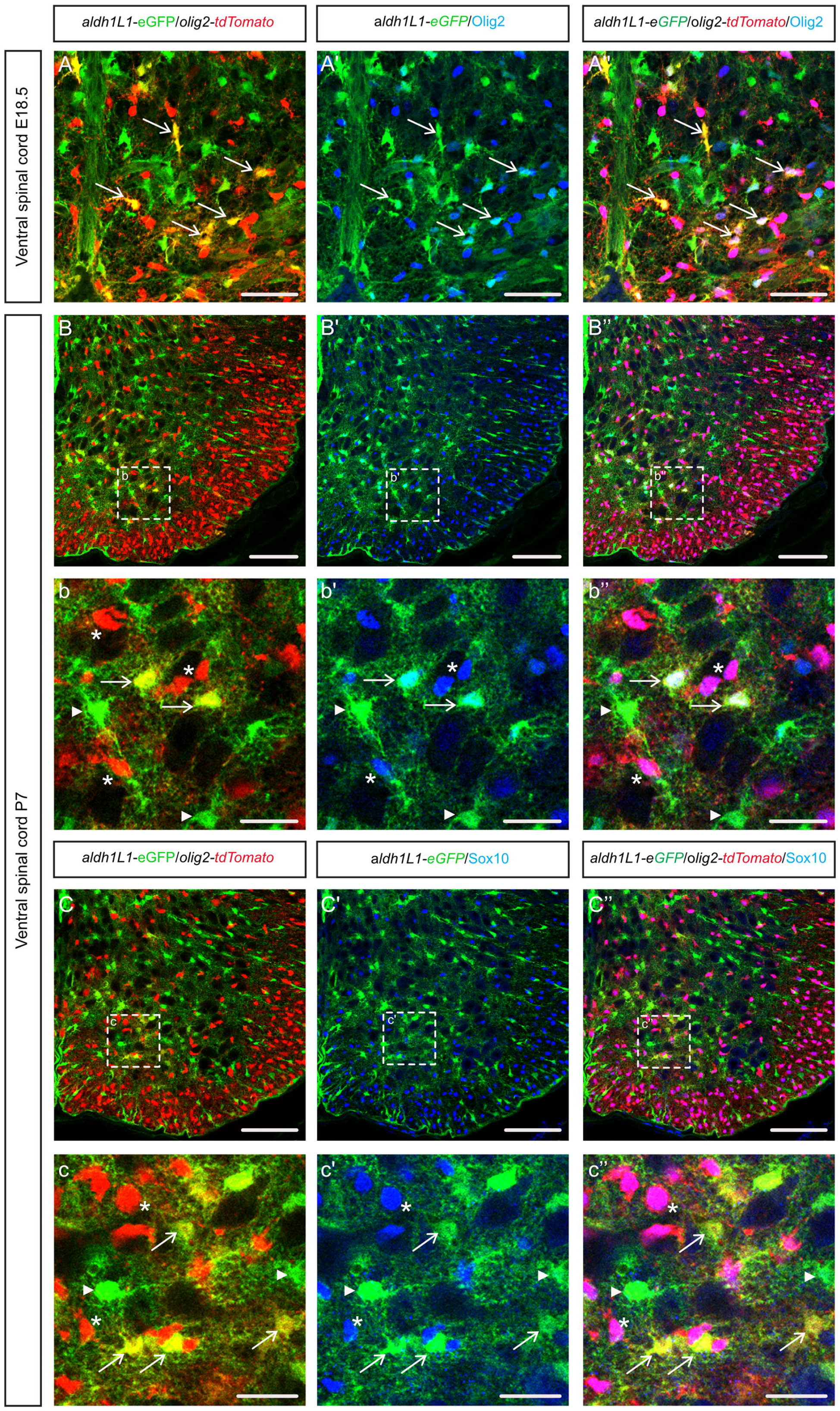
Detection of Olig2-AS in *aldh1L1-*GFP*/olig2*-tdTomato double transgenic mice. Here and in all subsequent panels, images show transverse sections of hemi-ventral spinal cord. **A-c’’**: Combined detection of GFP (green), td-Tomato (red) and Olig2 (blue, A-b’’) or Sox10 (blue, C-c’’) on *aldhlLl*-GFP/*olig2*-tdTomato transgenic mice at E18.5 (A-A”) and P7 (B-c”). Horizontal sets present successively GFP and tdTomato staining, Olig2 or Sox10 and GFP staining (prime) and the merged image (double prime). Images b-b” and c-c” show higher magnification of the areas framed in B and C, respectively. Asterisks, arrows and arrowheads point to OPC/OL, Olig2-AS and nonOlig2-AS, respectively. Scale bars = 100 μm in B-B”and C-C”; 50 μm in A-A” and 25 μm in b-b” and c-c”.

Overall, these data validate the *olig2-*tdTomato/*aldh1L1-*eGFP double transgenic mouse line as a suitable model to specifically color code the Olig2-AS and distinguish these cells from nonOlig2-AS and OPC/OL in the developing and post-natal spinal cords.

### Efficient RNA-seq based segregation analysis of Olig2-AS, nonOlig2-AS and OPC/OL

A first step towards the identification of differentially expressed genes between Olig2-AS and nonOlig2-AS or OPC/OL was to purify the three differentially labelled cell populations. For this purpose, spinal cord cells were dissociated at P7 (Figure S2A), a stage which has proven to be suitable for the purification of astrocytes with minimal activation and when, even if astrocytes are not fully differentiated, their gene expression profiles closely resemble that of mature astrocytes (Bushong et al., 2004; Cahoy et al., 2008). As a first approach, spinal cords from *aldh1L1-*eGFP and *olig2-*tdTomato simple transgenic mice were processed into single cell suspensions and sorted by FACS relying on either eGFP or tdT expression. Gating was then performed on these samples and further applied to all experiments (Figure 2A, B). We next FACS-sorted cells dissociated from spinal cords of *αldh1L1-*eGFP/*olig2*-tdTomato double transgenic mice (Figure 2C). This procedure was reproduced four times, each time with a different mouse litter, to ultimately get four replicates of tdT+, eGFP+ and eGFP+/tdT+ cell type populations. We extracted RNA from purified cell populations and performed RNA-Seq. The transcriptional profiles of each cell type replicate were then compared. To assess the reproducibility of our data and conservation across biological replicates, we calculated correlations across all RNA-Seq samples and indeed found high correlations among cell type replicates (Correlation coefficient R>0.92, Figure S2B). Principal-component analysis and hierarchical clustering data further showed good discrimination between the different cell type replicates according to mRNA expression profiles (Figure S2C-D). Our results, available online as a resource through GEO accession number GSE158517 (https://www.ncbi.nlm.nih.gov/geo/query/acc.cgi?acc=GSE158517), indicated that gene expression was similar in each group, thus providing a dataset we can use to find out the molecular specificity of the distinct glial cell populations.

**Figure 2:**
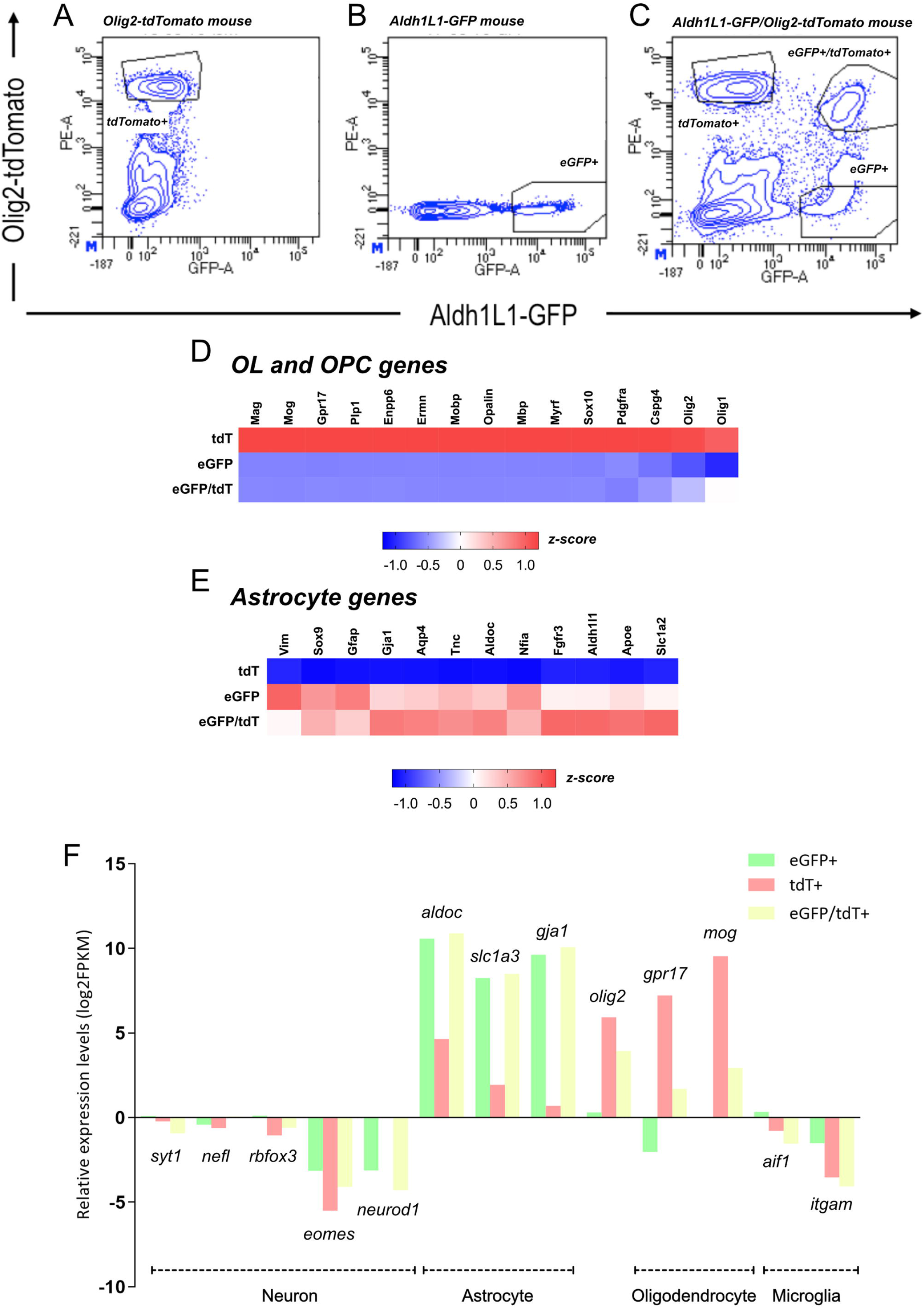
FACS purification and transcriptome analysis of eGFP+, tdT+ and eGFP+/tdT+ cell populations. **A-B**: Representative FACS plots showing the gating strategy. Gates for tdT+ (A) and eGFP+ (B) cells were defined from cell populations purified from *olig2*-tdTomato and *aldh1l1*-eGFP single transgenic mice, respectively. **C**: Representative FACS plot of the purification of tdT+, eGFP+ and tdT+/eGFP+ cell populations from P7 *aldh1l1*-eGFP/*olig2*-tdTomato double transgenic mice. **D-E**: Heatmap profiles of established oligodendrocyte and astrocyte specific genes in tdT+, eGFP+ and eGFP+/tdT+ FACS-sorted cell populations. **F**: Comparison of neuronal, astroglial, oligodendroglial and microglial gene expression levels (by log2FPKM values) in the three cell populations. Average FPKM values of four biological replicates are shown. All heatmaps are presented as z-score.

To validate the purity of the isolated cell types, we next probed the transcriptome data for the expression of eGFP and tdTomato for each cell type replicate. In agreement with the expected expression profiles for the two reporter proteins in each cell type replicate, we found high expression levels of eGFP in cells sorted on eGFP or eGFP and tdT expression and of tdTomato in cells sorted on tdT or tdT and eGFP expression (Figure S2E). We then, probed the transcriptome data for expression of well-known cell type specific genes for astrocytes, oligodendroglial cells, neurons and microglia (Cahoy et al., 2008; Zhang et al., 2014). Classic macroglial cell specific markers exhibited high expression levels in their corresponding cell types: the tdT+ cell population was enriched for OPC and OL genes while the eGFP+ and eGFP+/tdT+ cell populations were enriched for astrocyte genes (Figure 2 D-E). FPKM values of astrocyte genes, such as *aldoC, gjal* and *slcla3* were generally high in the eGFP+ and the eGFP+/tdT+ cell populations while oligodendroglial genes, such as *mog* and *gpr17,* displayed a high FPKM value in the tdT+ cell population (log2FPKM>7, Table S1, Figure 2F) but were undetectable or at extremely low expression levels by the remaining cell populations (log2 FPKM<2, Table S1, Figure 2F). Expectedly, expression of *olig2* was high in tdT+ and eGFP/tdT+ cells but quite undetectable in eGFP+ cells. By contrast, FPKM values of neuronal (e.g. *sytl, nefl, neurod1, rbfox3, eomes*) and microglial (e.g. *aifl* and *itgam*) specific genes were found negative (log2 FPKM<0, Table S1) in all the three cell populations (Figure 2F).

Together, these data confirmed the purity of the various isolated cell types and established the feasibility of constructing a high-quality transcriptome database representative of Olig2-AS (eGFP+/tdT+), nonOlig2-AS (eGFP+) and OPC/OL (tdT+).

### A specific Olig2-AS molecular signature revealed by RNA-Seq analysis

To identify an expression profile that distinguishes Olig2-AS from the other glial cell types, we carried out a two-step comparative analysis. We first determined the number of genes that are significantly upregulated (log2FoldChange (log2FC)>0 and p<0.05) or downregulated (log2FC<0 and p<0.05) in each cell population. For this purpose, we performed a two-by-two comparison by developing a pairwise differential analysis of RNA-seq using DESeq2 with normalized counts (Love et al., 2014). We thus ultimately got three differential analysis (DA), between Olig2-AS and nonOlig2-AS (DA1), between Olig2-AS and OPC/OL (DA2) and between nonOlig2-AS and OPC/OL (DA3) (Figure 3A). DA1 comparison showed a relatively low number of genes displaying differential values as only 813 and 1033 genes were found up-regulated and down-regulated, respectively (Figure 3A and S3A). In contrast, a large number of genes showed significant differential expression values in DA2 (7380 up-regulated and 5322 down-regulated genes) and DA3 (9094 up-regulated and 5885 down-regulated genes) in which Olig2-AS and nonOlig2-AS were compared to OPC/OL, respectively (Figure 3A and S3B-C). Thus, and as expected, these data indicated that Olig2-AS and nonOlig2-AS share many similarities but largely differ from OPC/OL. To identify an expression profile that distinguishes the Olig2-AS from the two other glial cell types, we further performed a second selection process aimed at identifying genes up-regulated in both DA1 (Olig2-AS versus nonOlig2-AS) and DA2 (Olig2-AS versus OPC/OL). In this process, the threshold was adjusted to log2FC>1 and padj<0.05. Analysis of the relative expression values then revealed a set of 135 highly expressed genes (Figure 3B, C, F and Table S2). We considered these genes as ‘signature’ genes as they together provide a unique profile of the Olig2-AS. Notably, among these genes, 39 were found up-regulated in the Olig2-AS but not in the nonOlig2-AS compared to OPC/OL (compare up1up2 and up3 in Figure 3A and S4A), indicating that these genes are good candidates to represent specific molecular markers of the Olig2-AS. Within this short list, only few genes, such as *ptn, grm5* or *lrtm2,* are recognized as astroglial genes (Gonzalez-Castillo et al., 2015; Sun et al., 2013; Chaboub et al., 2016) while the majority had not previously been identified as cell type specific.

**Figure 3:**
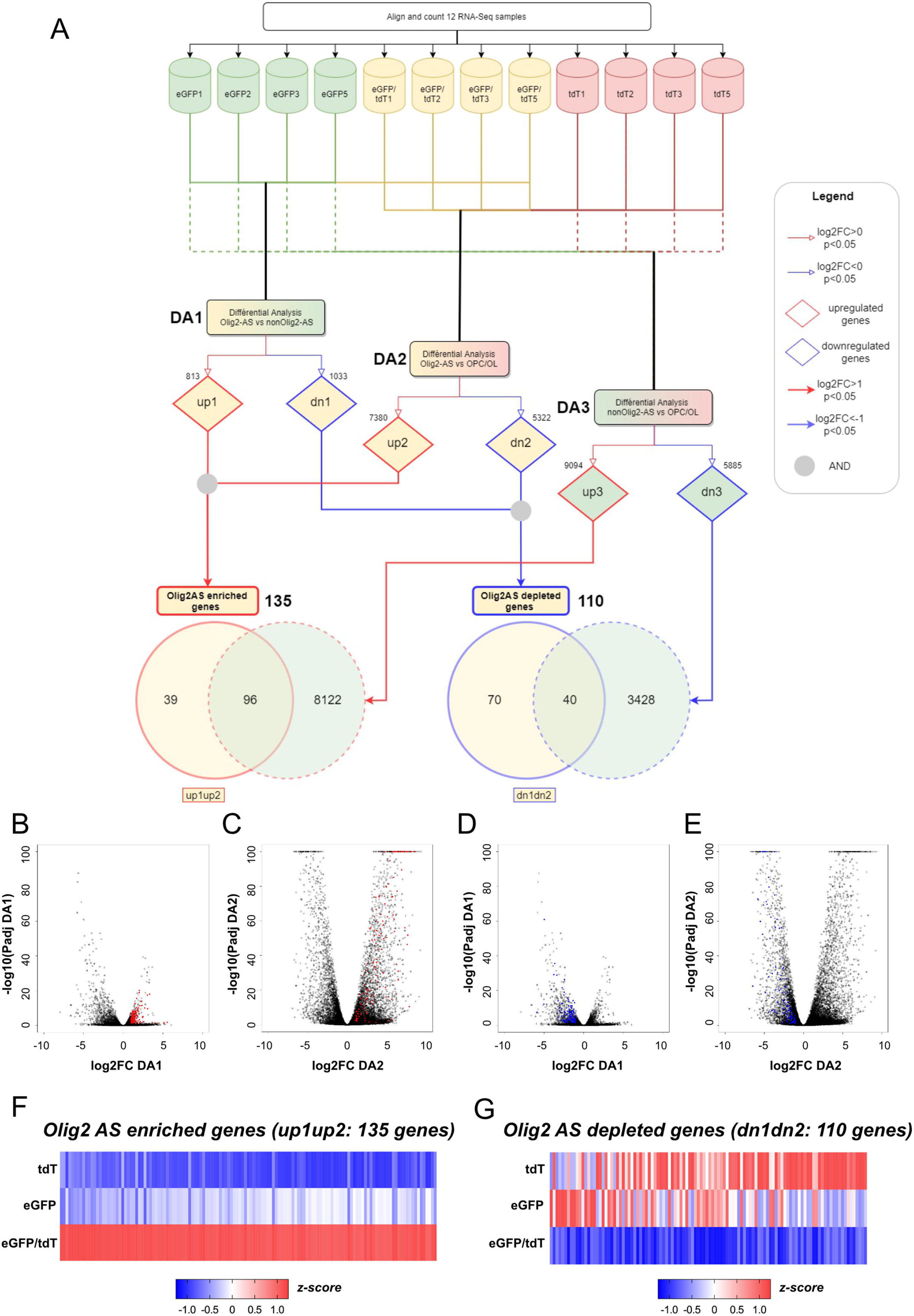
Identification of Olig2-AS molecular signature by differential analysis. **A**: Flowchart representing the treatment and two-step comparative analyses performed on the RNA-seq data. The top half of the chart shows the pooling of the 4 replicates for each cell population. The bottom half represent the two-consecutive comparisons between the glial cell subtypes. **B-E**: Volcano plots of differentially expressed genes representing the log2FC and -log10Padj from the differential analysis DA1 (Olig2-AS versus nonOlig2-AS, B, C) and DA2 (Olig2-AS versus OPC/OL, D, E). Red dots in B and C represent transcripts enriched in the Olig2-AS (up1up2; padj<0.05 and a log2FC>1). Blue dots in D and E represent transcripts down-regulated in Olig2-AS (dn1dn2, padj<0.05 and a log2FC<1). **F-G**: Heatmap analysis representing the 135 genes up-regulated (F) and the 110 genes down-regulated in Olig2-AS. All heatmaps are presented as z-score.

A similar analysis was performed for the genes down-regulated specifically in the Olig2-AS, adjusting the threshold to log2FC<-1 and padj<0.05. This yielded a list of 110 genes down-regulated both in DA1 and DA2 (Figure 3D, E, G and Table S3). As noted above, among these genes, 70 were found down-regulated in the Olig2-AS but not in the nonOlig2-AS compared the OPC/OL (Figure 3A and S5A). This analysis thus revealed a set of genes whose expression is repressed specifically in the Olig2-AS subtype, thus reinforcing the view that the molecular identity of Olig2-AS differs from that of nonOlig2-AS.

### Validation of RNA-Seq results by in situ hybridization and identification of Olig2AS specific markers

The bioinformatics analysis has generated a list of 135 genes enriched in the Olig2-AS, including 39 potential molecular markers of Olig2-AS. For validation, we selected, from this short list, two genes, *inka2* and *kcnip3,* whose expression in astrocytes had not been previously reported and performed their expression pattern analysis on P7 spinal cord sections. Both *inka2* and *kcnip3* mRNA were detected in cells scattered in the ventral spinal grey matter while only few positive cells were found in the white matter, thus revealing cells displaying a distribution very reminiscent to that of Olig2-AS (Figure 4A, B). To confirm that these cells indeed are Olig2-AS, we next combined detection of *inka2* or *kcnip3*mRNAs with immunodetection of Olig2 and in situ localization of the *fgfr3* mRNA, an early and specific hallmark of astroglial cells (Pringle et al., 2003; Ohayon et al., 2019). We found that most, if not all, *inka2+* and *kcnip3+* cells were positive for both Olig2 and *fgfr3* stainings (Figure 4C-D’’’), thus validating *inka2* and *kcnip3* as new specific markers of Olig2-AS. Because *inka2* has previously been reported to be an OPC marker in the developing spinal cord (Iwasaki et al., 2015), we examined its expression, together with that of Olig2 and *fgfr3,* in the E14.5 embryonic spinal cord. As observed at P7, we found that *inka2* positive cells all express the *fgfr3* mRNA (Figure S6A, B) and comparison with the Olig2 staining showed *inka2+/fgfr3+* cells positive for the Olig2 staining intermingled with *inka2+/fgfr3+* cells that do not express Olig2 (Figure S6A’-B’’). In these experiments, we never found cells co-expressing *inka2* and Olig2 but not *fgfr3.* These data thus indicate that, during development, expression of *inka2* marks astrocyte precursor cells but not OPC and that, unlike at post-natal stage P7, *inka2* expression is not restricted to the Olig2-AS sub-type but also marks at least a subset of nonOlig2-AS precursor cells. We thus conclude that the preferential expression of *inka2* in Olig2-AS observed at P7 results from down-regulation of this gene during the maturation period of nonOlig2-AS.

**Figure 4:**
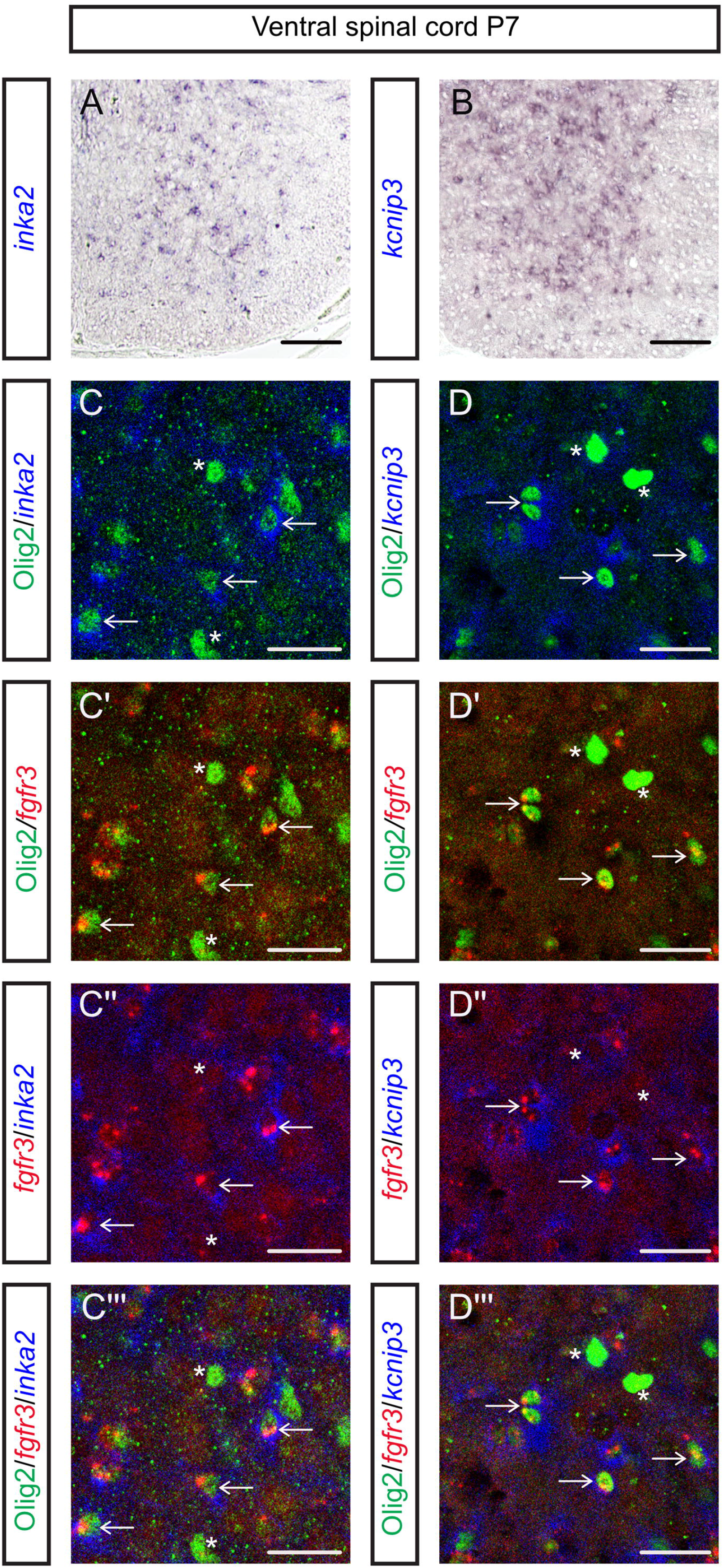
The up-regulated genes *inka2* and *kcnip3* are specifically expressed in Olig2-AS. **A-B:**Expression profiles of *inka2* (A) and *kcnip3* (B) mRNAs at P7. **C-D’’’**: Combined detection of Olig2 (green),*fgfr3* (red) and *inka2* (blue, C-C”‘) or *kcnip3* (blue, D-D’”) mRNAs. Vertical sets present successively Olig2 and *inka2*or *kcnip3* mRNA detection (C, D), Olig2 and*fgfr3* staining (C’, D’), *fgfr3*and *inka2* staining (C’’, D’’) and the merged image (C’’’, D’’’). Asterisks and arrows point to OPC/OL and Olig2-AS, respectively. Scale bars = 100 μm in A and B and 25 μm in C-C’’’ and D-D’’’.

Still focused on validating the RNA-seq, we selected one gene among the 70 genes found down-regulated in Olig2-AS (Figure S5A). The selected gene was *efnb3,* a member of the *ephrin* gene family, previously reported as expressed in postnatal myelinating OLs of the mouse spinal cord (Benson et al., 2005). Accordingly, we preferentially detected the *efnb3* mRNA in cells located in the white matter of the P7 spinal cord (Figure S7A). Detection of *efnb3* mRNA was next combined with immunodetection of Olig2 and in situ localization of the *fgfr3* mRNA. We found no *fgfr3*+/Olig2+ cell positive for the *efnb3* mRNA staining in the grey matter (Figure S7C-C’’) but, as expected, we detected numerous *efnb3+/Olig2+* cells negative for the *fgfr3* staining in the white matter (Figure S7D-D’’). These data thus reinforce the conclusion that the RNA-seq approach reliably identifies gene downregulated in Olig2-AS.

### Functional analysis of genes differentially expressed in Olig2-AS

To get insights on the biological meaning behind the list of the 135 and 110 genes enriched and decreased in Olig2-AS, respectively, we performed Gene Ontology (GO) term-enrichment analysis (GO summaries). In terms of biological processes, Olig2-AS enriched genes were classified into 5 functional groups: “central nervous system development”, “chemical synaptic transmission”, “carboxylic acid transport”, “regulation of synapse organization” and “response to steroid hormone” (Figure 5A, Table 1, Table S4). Among these 5 GO-terms, 3, including “chemical synaptic transmission”, “carboxylic acid transport” and “regulation of synapse organization”, are related to the synapse (Figure 5A). On the other hand, GO analysis performed on down-regulated genes displays a strong association with “regulation of cell development”, including “neuron projection development” (Figure 5B, Table1, Table S4). To get more details, circular visualization of the GO results was performed selecting synaptic-related and cell development-related terms (Figure 5C). Data indicated that, among the 135 genes enriched in Olig2-AS, 16 are associated with GO synapse-related terms while only 2 among the 110 down-regulated genes are associated with these terms (Figure 5C, Table1). Instead, down-regulated genes appear mostly associated to cell development terms (Figure 5C, Table1). Thus, GO annotation results pointed to 16 genes which are dominant in Olig2-AS and are related to modulation of chemical synaptic transmission and glutamate receptor signaling pathways. Finally, GO analysis of genes up-regulated in Olig2-AS performed in terms of molecular function (MF) again pointed to synaptic related functions such as “carboxylic acid transmembrane transporter activity”, including “active transmembrane transporter activity” and “sulfur compound binding” (Figure 5D). Among these genes, we identified genes coding for the Glutamate Metabotropic Receptors 3 and 5 (*grm3* and *grm5*), the GABA receptor subunits (*gabbr1* and *gabbr2*), the glutamine transporter SNAT3 (*slc38a3*) and the glycine and D-serine transporter SLC7A10 (*slc7a10*). To go further and have a broader view of the molecular identity of Olig2-AS, we next compared expression levels in Olig2-AS and nonOlig2-AS of a set of genes encoding for neurotransmitter receptors and transporters (Figure 5E). We found that 3 (*grm3/5/7*) out of the 8 genes coding for glutamate metabotropic receptors are expressed in both Olig2-AS and nonOlig2-AS with significant enrichment for *grm3* and *grm5* in Olig2-AS (Figure 5E). Data indicated that Olig2-AS and nonOlig2-AS also differ by *grm8* expression which appears significantly enriched in the nonOlig2-AS (Figure 5E). Similar analysis performed for genes coding for ionotropic glutamate receptors indicated that only 2 genes encoding for NMDA subunits (*grin2b*, *grin3a*) are expressed in spinal astrocytes with no difference in their expression levels (Figure 5E). All out of the 4 genes encoding for AMPA receptors (*gria1-4*) were found expressed in spinal astrocytes with no significant difference in their expression levels except for *gria1* which is specifically enriched in the nonOlig2-AS (Figure 5E). We also analysed genes encoding for AMPA receptor interacting proteins which are known to modulate localization and functional properties of the receptors (Jacobi and von Engelhardt, 2018; Molders et al., 2018) and found enrichment for 3 of them (*gsg1l, shisa6* and *shisa9*)in Olig2-AS compared to nonOlig2-AS (Figure 5F). We further analysed a set of genes whose products are known to control glutamate homeostasis, i.e. the glutamate transporters GLT1 (*slc1a2*) and GLAST-1 (*slcla3*), the glutamine synthetase GLUL (*glul*), the vesicular glutamate transporters VGLUT1 (*slc17a7*) and VGLUT2 (*slc17a6*) and the sodium-dependent glutamine transporters SN2 (*slc38a5*) and SN1 (*slc38a3*), the latter being the only gene found enriched in Olig2-As compared to nonOlig2-AS (Figure 5G). We also found expression of a majority of genes known to encode for different subunits of GABA receptors both in Olig2-AS and nonOlig2-AS (Figure 5H). Noticeably, 2 genes, encoding for the metabotropic (*gabbr2*) and ionotropic (*gabrg1*) GABA receptors, were found enriched in Olig2-AS (Figure 5H). Three GABA transporters GAT1/3/4 (*slc6al, slc6a13, slc6all*) as well as 2 glycine transporters (*slc6a9* and *slc7alO*) were also found expressed in the two astrocyte subsets with significant enrichment in Olig2-AS of *slc6al, slc6all,* coding for GABA transporters, and of *slc6a9* and *slc7alO,*coding for glycine transporters (Figure 5I). Together, these analyses, pointing to differential expression in Olig2-AS and nonOlig2-AS of genes involved in synaptic transmission with a set of 12 genes (*grm3, grm5, gsg1l, shisa6, shisa9, slc38a3, gabbr2, gabrg1, slc6al, slc6all, slc6a9* and *slc7alO*) enriched in Olig2-AS and 3 genes (*grm8, gria1* and *slc6a13*) conversely enriched in nonOlig2-AS, provide clues as to how the Olig2AS might differ in their ability to dynamically modulate synapse function.

**Table 1:**
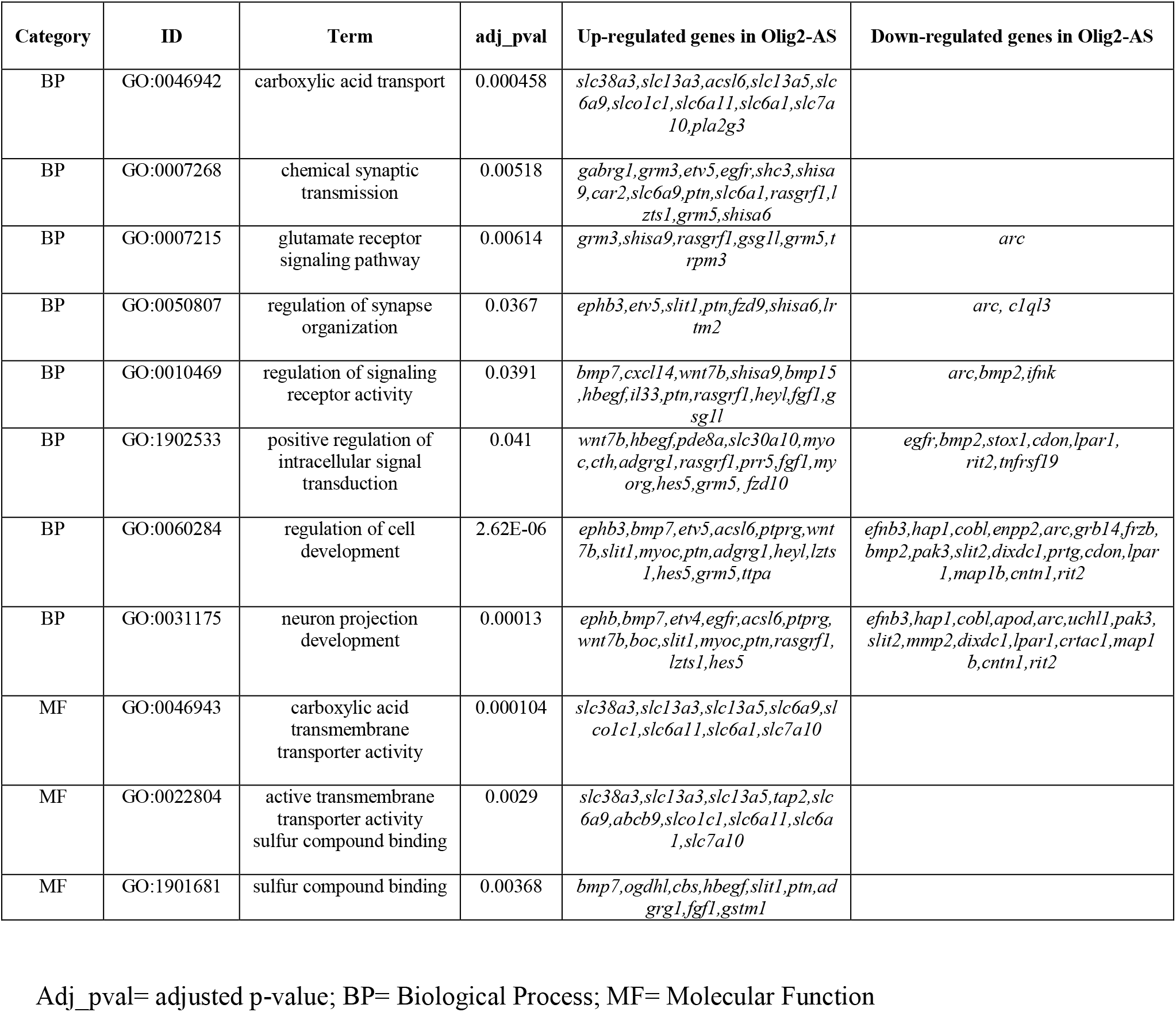
Gene ontology of all 245 differentially expressed genes in Olig2-AS.

**Figure 5:**
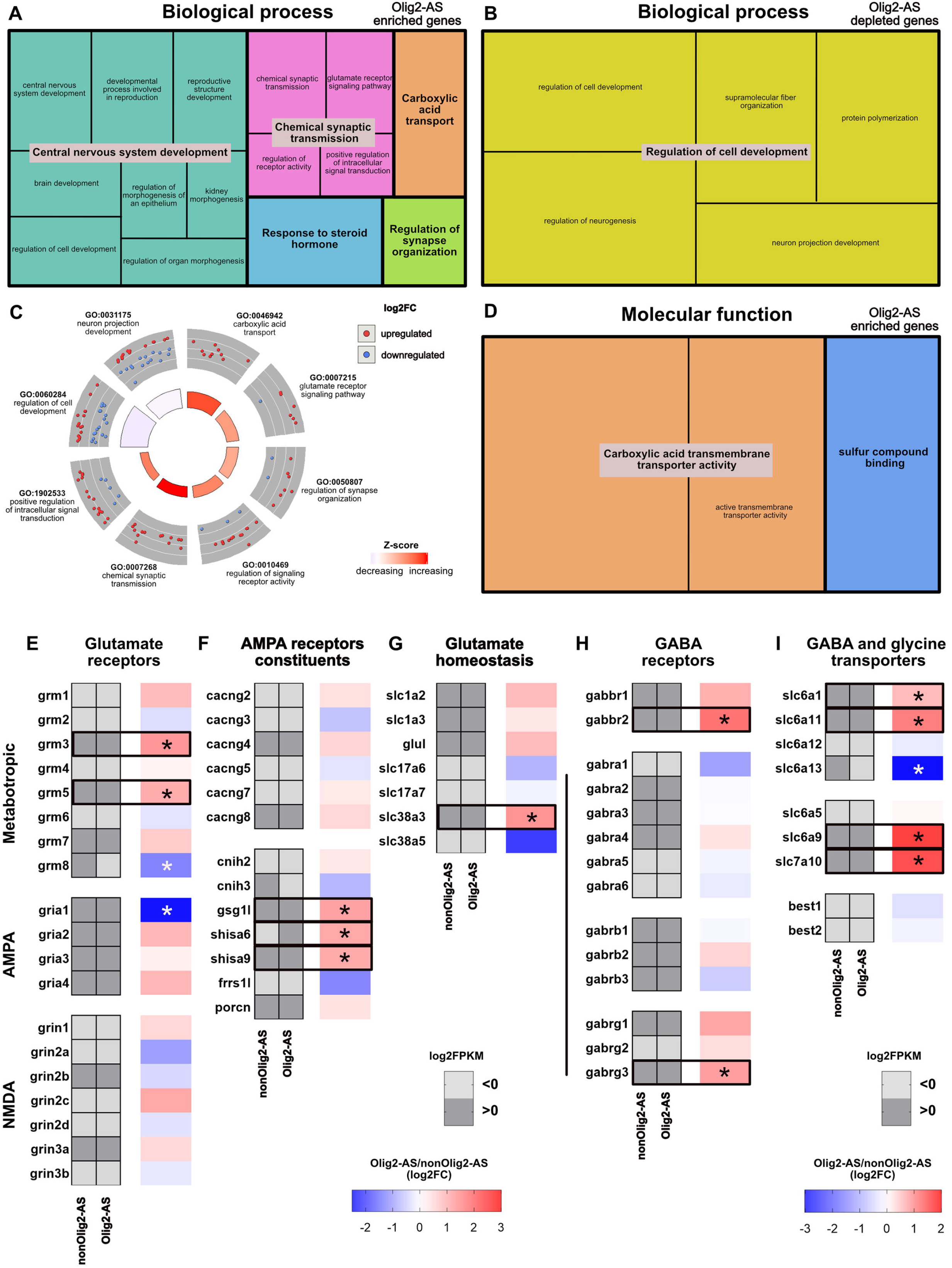
Functional enrichment analysis of Olig2-AS differentially expressed genes. **A-D**: Treemap representations of Gene Ontology (GO) (A, B and D) and circular visualization of geneannotation enrichment (GOcircle, C) analyses performed on Olig2-AS differentially expressed genes. GO analyses were performed from genes specifically up-regulated (A, D) or down-regulated (B) in Olig2-AS and are presented in terms of Biological Process (A, B) and Molecular Function (D). Significant GO terms (smaller squares) were grouped according to their parental ontology to highlight highly represented functions. Inner circle in C depicts the main biological processes found increased (red) or decreased (blue) in Olig2-AS. The outer circle shows scaled scatter plots for involved genes and their regulation within the most-enriched biological processes (GO term). Blue dots mark downregulated, and red dots mark upregulated genes in Olig2-AS (log2FC). The z-score (inner circle) defines the likelihood of a process being decreased (blue) or increased (red) in Olig2-AS. **E-I**: Heatmaps depicting abundance of mRNA encoding for glutamate receptors (E), AMPA receptors constituents (F), glutamate transporters and enzyme (G), GABA receptors (H) and GABA and glycine transporters (I). Heat maps in gray represent mRNA expression levels (log2FPKM) in nonOlig2-AS and Olig2-AS. Light and dark grays indicate values lower and above zero, respectively. Adjacent heatmaps illustrate fold-change in abundance of mRNA in Olig2-AS versus nonOlig2-AS (log2FC). Black and white asterisks indicate significant upregulation and downregulation in Olig2-AS, respectively.

### Additional functional (dis)similarities between Olig2-AS and nonOlig2-AS

Beyond their function in regulating synaptic activity, astrocytes are also known to carry out important functions in synapse formation and elimination (Chung et al., 2015) and in regulating homeostasis and energetic support (Sofroniew and Vinters, 2010). They are also known to be a primary source of lipid synthesis in the brain and, in particular, to provide neurons with cholesterol which is essential for presynaptic vesicle formation (Kiray et al., 2016). None of these functions were found explicitly over- or under-represented in the above-mentioned GO term-enrichment analysis. However and in particular because this analysis was performed from a relatively limited list of genes, we examined the possibility that Olig2-AS and nonOlig2-AS might differ by expression levels of genes involved in these functions. We first analysed expression levels of genes coding for key players of synapse formation, i.e. Thrombospondins (*thbs 1, 2 and 4*), SPARC-like protein 1/Hevin (*sparcl1*), SPARC (*sparc*)(Christopherson et al., 2005; Kucukdereli et al., 2011), Neuroligines (*nlgn1-3*), Neurexines (*nrxn1-3*)(Stogsdill et al., 2017; Uchigashima et al., 2019) and Glypicans (*gpc4, gpc6*) (Allen et al., 2012), or synapse elimination, i.e. members of the Transforming growth factor beta family (*tgfb1-3*), the tyrosine kinase receptor MERTK (*mertk*) and the protein MEGF10 (*megf10*) (Chung et al., 2013). All of these genes, except *thbs4* and *tgfb1,* were found expressed in Olig2-AS and nonOlig2-AS with no significant difference in their expression levels, if not a trend towards down-regulation of the 3 *thbs* genes in the Olig2-AS (Figure 6A). Similar analyses were then performed for a set of genes whose products are known to control brain homeostasis and energetic support, i.e. the Aquaporin-4 water channel (*aqp4*),the inwardly rectifying potassium channels Kir4.1 (*kcnj10*) and Kir5.1 (*kcnj16*) (Simard and Nedergaard, 2004), the glucose transporter GLUT1 (*slc2a1*) and components of the lactate shuttle (*slc6a1*, *slc16a3, ldha*) (Tekkok et al., 2005). Again, all of these genes, except *slc16a3,* were found expressed both in Olig2-AS and nonOlig2-AS with no significant difference in their expression levels (Figure 6B). Finally, we examined expression levels of a set of genes involved in cholesterol synthesis, i.e. different enzymes, HMGCR (*hmgcr*), DHCR7 (*dhcr7*), DHCR24 (*dhcr24*), FDFT1 *fdft1*), ACAT1 (*acat1*), ACAT2 (*acat2*) and CYP51 (*cyp51*) (Pfrieger and Ungerer, 2011) and the two transcription factors regulating lipid synthesis, Srebf1 (*srebf1*) and Srebf2 (*srebf2*). All of these genes were expressed in Olig2-AS and nonOlig2-AS and no significant difference were found in their expression levels (Figure 6C). However, some differences can be observed when looking at the expression of the cholesterol binding molecules Apolipoprotein D (*apod*), Apolipoprotein E (*apoe*) and Apolipoprotein J (*clu*) and of specific transporters, i.e. ATP binding cassette (ABC) transporters *abcg1, abcg4* and *abca1* (Pfrieger and Ungerer, 2011). Among genes found expressed by both Olig2-AS and nonOlig2-AS, 2, *abcg1* and *apod,* appeared specifically down-regulated in Olig2-AS (Figure 6C). Taken together, these analyses did not reveal major differences between Olig2-AS and nonOlig2-AS in relation to these important physiological functions of astrocytes.

**Figure 6:**
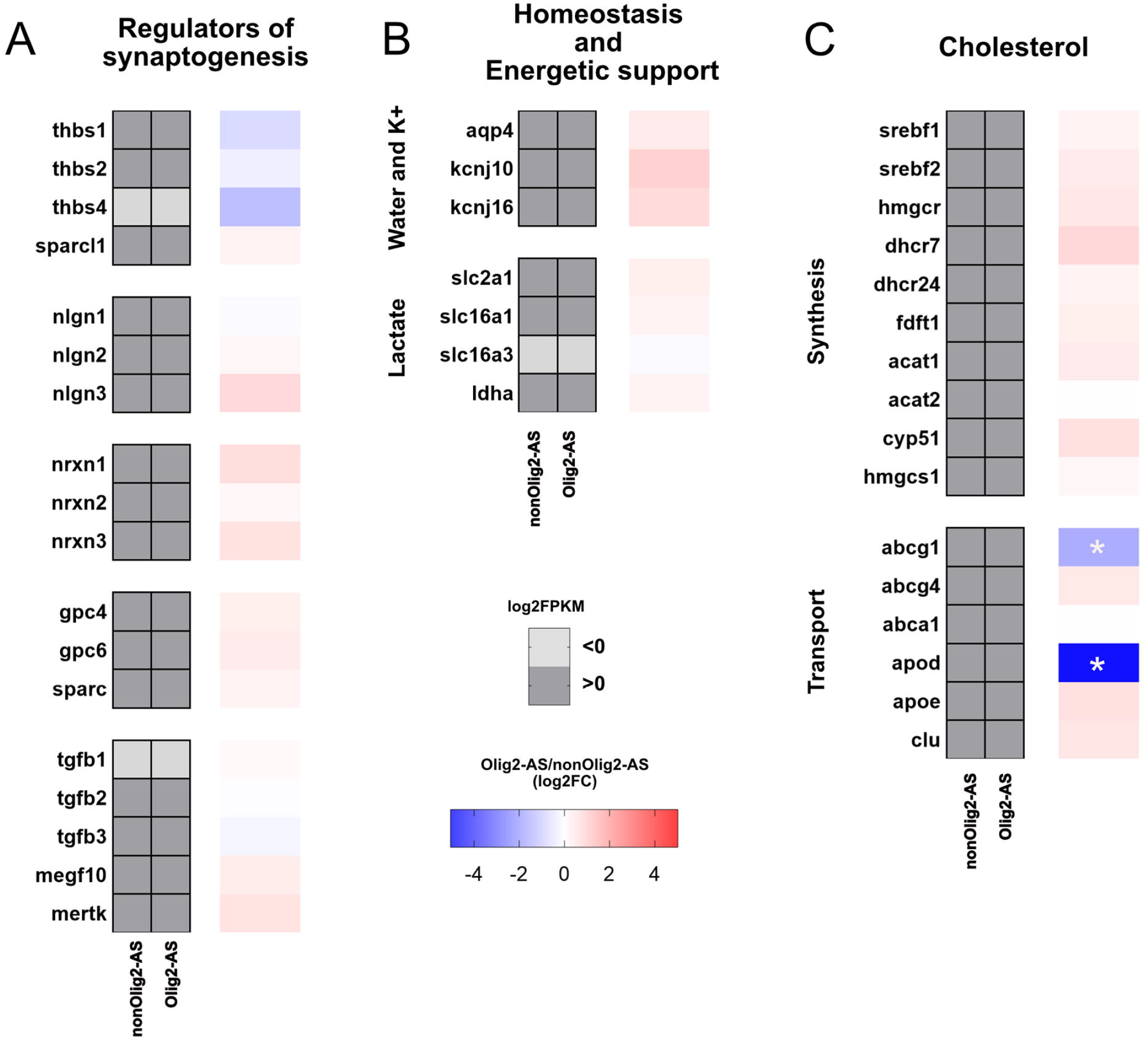
Functional similarities between Olig2-AS and nonOlig2-AS. **A-C**: Heatmaps depicting abundance of mRNA encoding for genes involved in synaptogenesis (A), homeostasis and energetic support (B) and cholesterol metabolism (C). Heat maps in gray represent mRNA expression levels (log2FPKM) in nonOlig2-AS and Olig2-AS. Light and dark grays indicate values lower and above zero, respectively. Adjacent heatmaps illustrate fold-change in abundance of mRNA in Olig2-AS versus nonOlig2-AS (log2FC). White asterisks indicate significant downregulation in Olig2-AS.

## Discussion

In recent years, transcriptome profiling has been widely used to unravel astrocyte heterogeneity and a large amount of data supporting molecular astrocyte diversity has been provided. However, how this diversity is defined, i.e. does it result from specific developmental transcriptional programs and/or from later refinement by local environmental cues remains an open question. In this context, the main novelty of our work is that it allows performing relationship between molecular identity of a particular astrocyte subset and its developmental origin. The method we set up made it possible to isolate the Olig2-AS a sub-population of spinal cord astrocytes which originate from Olig2/pMN progenitors (Ohayon et al., 2019), from the other spinal glial cell populations, i.e. OPC/OL, also generated by Olig2/pMN progenitors, and other astrocyte subsets that do not express Olig2 (nonOlig2-AS) and are generated by more dorsal progenitors which express patterning factors distinct from Olig2 (Bayraktar et al., 2015). Comparison of RNA-Seq transcriptome datasets obtained for the three cell types revealed a unique Olig2-AS molecular profile, thus providing, for the first time, a specific gene expression signature of an astrocyte subtype whose identity is tied to its embryonic origin.

A major limitation in understanding astrocyte diversity is the lack of specific markers to prospectively identify and isolate the distinct astrocyte subtypes. A first important finding of our work is thus the identification of two highly specific expression markers of Olig2-AS in the post-natal spinal cord, *kcnip3* and *inka2*, both entering the list of genes composing the Olig2-AS molecular signature. *Kcnip3* encodes for the voltage-dependent K+ channel interacting protein 3 (KNIP3), also known as Calsenilin/KChIP2/DREAM. This multifunctional protein is known to interact with the Kv4 channels and modulate A-type potassium currents in a Ca2+-dependent manner but also to function as a calcium-regulated transcriptional repressor (An et al., 2000; Ledo et al., 2000; Anderson et al., 2010). The second gene, *inka2*, encodes for the Inka Box Actin Regulator 2, an inhibitor of the serine/threonine-protein kinase PAK4 (Iwasaki et al., 2015). Expression of *inka2* in Olig2 positive cells of the developing spinal cord has previously been reported and this observation led, at this moment, to the conclusion that expression of this gene marks cells of the oligodendroglial lineage (Iwasaki et al., 2015). Our present data show that co-expression of *inka2* and Olig2 instead marks the Olig2-AS which also expresses Olig2 from the earliest stage of their generation in the ventral spinal cord (Ohayon et al., 2019).

Additional exploration into the molecular signature of the Olig2-AS points to a list of enriched genes that encode for secreted factors, transcriptional regulators and factors associated with synapse function. First on this list is *ptn* encoding for Pleiotrophin (PTN), a secreted growth factor with multiple functions during neural development, also known to be up-regulated in astrocytes in response to injury and to act as a neurotrophic factor for spinal motor neurons (Mi et al., 2007; González-Castillo et al., 2015). Another important result of our work is that we identify a number of transcriptional regulators (ERRα, Zfp219, Kcnip3, Irx2, Etv5, HES5, Etv4, Heyl, Epas-1), all likely contributing to the control of the unique expression profile observed in Olig2-AS. To our knowledge, most of them have not previously been linked with astrocyte identity or function. It may be noted that HES5 is known to enhance astrogliogenesis (Bansod et al., 2017) and that Etv5 (ERM) has recently been reported to be differentially expressed in cortical astrocytes (Batiuk et al., 2020). Finally, the most noteworthy aspect is enrichment of a set of genes encoding for neurotransmitter receptors and transporters, mainly related to the glutamatergic and GABAergic signalling, thus supporting the view that Olig2-AS are endowed with particular functions in regulating synaptic activity in the ventral grey matter of the spinal cord. The proposal that local ventral horn astrocytes might ensure specialized function is not a new idea. Previous work from D. Rowitch and collaborators has indeed highlighted the presence, in this region, of a specialized astrocyte subset which plays a critical role to selectively maintain the physiological properties and function of MN subtypes (Molofsky et al., 2014; Kelley et al., 2018). Of note, these astrocytes were found enriched in transcripts for *kcnjlO,* which encode for the K+ channel Kir4.1 (Kelley et al., 2018). Although *kcnjlO* was not listed in the Olig2-AS molecular signature, this gene was found enriched with a fold change value of 0.98 compared to nonOlig2-AS, i.e. a value just below the arbitrary fold change cut-offs of >1 that we have chosen to apply. This observation, together with the close association of Olig2-AS and MNs (Ohayon et al., 2019), leaves open the possibility that the Olig2-AS correspond to those astrocytes identified for their discrete molecular function in supporting alpha MN physiology. Otherwise, it is also interesting to note the recent identification of a regionally and functionally distinct astrocyte subset, located in cortical layer V of the cerebral cortex (Miller et al., 2019), which presents notable similarities with Olig2-AS in the sense that they were found enriched in *olig2* and *lgr6* transcripts, the latter being one of the 135 genes enriched in Olig2-AS. These cortical astrocytes were also found enriched in *kcnjlO* and *ndp* (Norrin) transcripts, that, as previously mentioned for *kcnjlO,* were also found enriched in Olig2-AS but not included in the Olig2-AS molecular signature because of a fold change value below the cut-offs of >1. Although still very speculative, and because layer V of the cerebral cortex contains corticospinal neurons that regulate voluntary motor control (Anderson et al., 2010), it is tempting to hypothesize that Olig2 expression in astrocytes, either in the spinal cord or in the cerebral cortex, might confer them specific functional properties to control motor circuits.

In conclusion, our work provides the first transcriptomic study of an astrocyte subtype whose molecular identity can be linked to its developmental origin. We make available transcriptome dataset of this particular astrocyte subset (Olig2-AS) as well as that of other spinal cord astrocytes and oligodendroglial cells. Analysis of the 135 genes that compose the molecular signature of Olig2-AS reveals specific markers of this astrocyte subtype and provides a wealth of information to further unravel to what extent the developmental origin of an astrocyte subtype might influence its functional specialization.

## Authors’ contributions

DO, MA and NE performed and analysed experiments. DO and CS supervised the project and wrote the manuscript.

## Acknowledgements

We especially thank N. Rouach and P. Durbec for sharing of Aldh1L1-GFP and Olig2-tdTomato mice respectively, E. Nasser for the technical support at the IPBS-FACS platform. We acknowledge I. Néant, A. Davy and S. Sakakiba for the gift of the *kcnip3, efnb3* and *inka2* probes respectively and A. Davy, W. Richardson and JP Hugnot for critical reading of the manuscript. We thank the ABC facility and ANEXPLO for housing mice and the Toulouse Regional Imaging Platform (TRI) for technical assistance in confocal microscopy. We acknowledge the Developmental Studies Hybridoma Bank, developed under the auspice of NICHD and maintained by the University of Iowa, Department of Biological Sciences, Iowa City, IA, for supplying monoclonal antibodies. Work in C.S.’ lab was supported by grants from ANR, ARC (PJA20161204698), ARSEP, CNRS and University of Toulouse.

## Additional information

The authors declare that they have no competing interests.

## STAR Methods text

### RESOURCE AVAILABILITY

#### Lead contact and materials availability

Further information and requests for resources and reagents should be directed and will be fulfilled by the Lead Contact, David Ohayon (davi).

#### Data and Code Availability

The accession number for the RNA-Seq data reported in this paper is GEO: GSE158517. (https://www.ncbi.nlm.nih.gov/geo/query/acc.cgi?acc=GSE158517)

### EXPERIMENTAL MODEL AND SUBJECT DETAILS

#### Mouse strains

All procedures were performed in agreement with the European Community guiding principles on the care and use of animals (Scientific Procedures) Act, 1987 and approved by the national Animal Care and Ethics Committee (APAFIS#20396) following Directive 2010/63/EU. *Olig2-*tdTomato mice were generated by the Jackson lab (Tg(*olig2*-tdTomato)TH39Gsat) and were genotyped using the following primers (Tomato-Fw: 5’-CTGTTCCTGTACGGCATGG-3’ and Tomato-Rev: 5’-GGCATTAAAGCAGCGTATCC-3’). *Aldh1L1-*GFP (Cahoy et al., 2008) transgenic mice were genotyped as previously reported (Gong et al., 2003; Heintz, 2004). All mice were maintained on a 12h light/dark cycle with food and water ad libitum. Both males and females were used for all experiments. All mice were maintained on a C57BL/6 background.

### METHODS DETAILS

#### Dissociation procedure

Spinal cords from *aldh1L1-*GFP/*olig2*-tdTomato mice were harvested at postnatal day 7 (P7), only the brachial and thoracic levels were isolated. DRGs and meninges were then removed spinal cord explants were sliced into 100 μm sections using a tissue-chopper (Mc Ilwain). The tissue was enzymatically dissociated to make a suspension of single cells as described previously (Cahoy et al., 2008). Briefly, the tissues were dissociated with papain (20U/mL, Worthington Biochemical, CatN#LK003150) for 60 minutes at 37°C in bicarbonate-buffered Earle’s balanced salt solution and 0,005% of DNaseI (Worthington Biochemical, CatN#LK003150). The papain solution was equilibrated with 5% CO2 and 95% O2 before andduring papain treatment. Then, the tissue was mechanically dissociated by gentle trituration with a 10mL pipette. The cloudy cell suspension was collected and centrifuged at 300 x g for 5 minutes. The cell pellet was resuspended in a low concentration inhibitor solution with DNase, BSA 1mg/mL and ovomucoid 1mg/mL (Worthington Biochemical, CatN#LK003150). Then, the cell suspension was layered on top of 5mL of a high concentration inhibitor solution with BSA 10mg/mL and ovomucoid 10mg/mL and centrifuged at 70 x g for 6 minutes. The cell pellet was then resuspended in 300μL of PBS and 10% FBS (Gibco, CatN# A31604-01) and filtered through a 40μm cell strainer (Falcon, CatN#352340) to remove any remaining clumps of tissue.

#### FACS and RNA preparation

Dissociated cells from two mouse pups coming from the same litter were pooled. Cells were sorted using a FACSAria Fusion BSL1 at the IPBS FACS-platform. Dead cells and debris were gated by their low forward scatter area (FSC-A) and high side scatter area (SSC-A). Following that, two successive gating approaches have been done to exclude doublets, first a forward scatter height (FSC-H) vs forward scatter width (FSC-W) and then a side scatter height (SSC-H) vs side scatter width (SSC-W). Then a gating for fluorescence has been performed, the three awaited cells population were then identified based on their fluorescence using a PE-A vs GFP-A: high eGFP fluorescence for the nonOlig2-AS; high tdTomato fluorescence for the OPC/OL and high eGFP and tdTomato fluorescence for the Olig2-AS. Purified cells were harvested by centrifugation at 2000xg for 5 minutes and cell pellets were processed for RNA extraction. The whole procedure for cell suspension preparation and FACS sorting was completed in 3-4 hours. After sorting, RNA was isolated from the cell pellet using standard Trizol reagent (Thermofisher, Cat.N#15596026) with glycogen added as carrier according to manufacturer instructions and eluted into 35μL RNase-free water. Eluted RNAs was then stored at −80°C. RNA quality and integrity (RIN) were verified on an Agilent 2100 Bioanalyzer. The whole procedure going from the dissociation to RNA preparation has been performed four times, each time on a different mouse litter.

#### RNA-seq library generation and sequencing

For each cell population (eGFP+; tdT+; tdT/eGFP+) that was subjected to RNA-seq analysis, between 7×10^3^ and 2×10^4^ FACS isolated cells were used from pooled samples; each cell population was sequenced in quadriplicate. Library preparation and RNA sequencing on the twelve samples were performed by Beijing Genomics Institute (BGI, Hong Kong, China) usinga BGISEQ-500 RNA-seq platform (paired-end sequencing, 100 bp reads). The quality of each raw sequencing file (fastq) was verified with FastQC. All files were aligned to the reference mouse genome (Ensembl *mus musculus* GRCm38.91; gtf(mm10)) using STAR aligner (Version Galaxy Tool: V2.1; Dobin et al., 2012). Read count per sample was computed using HT-seq count (Version Galaxy Tool: V1.0; Anders et al., 2014). Then raw count table was_cleaned, we decided to keep gene having an averaged raw read count per sample higher or equal to 1.

#### Differential analysis

This analysis was applied per pairs of cell populations using DESeq2 (V1.22.2; Love et al., 2014), available as an R package in bioconductor (www.biocondutor.org), to normalize raw read count using RLE methods generating the log2 fold change (log2FC) values. This procedure has led to three different differential analysis (DA), between Olig2-AS and nonOlig2-AS cells (DA1), between Olig2-AS and OPC/OL (DA2) and between nonOlig2-AS and OPC/OL (DA3). Mouse ENSEMBL IDs from each sample were then converted to mouse official gene names using Biomart (Durinck et al., 2005), available also as an R package in bioconductor. We used principal component analysis (PCA) and hierarchical clustering to cluster samples based on their expression levels. Data representation of the PCA was generated using the ggplot2 package (https://ggplot2.tidyverse.org). Euclidean distances between the cell types were determined for the top 20% most variable mRNA and plotted using the heatmap.2 package in R software.

#### Sample correlation

Following RNA-seq data preprocessing, we computed the pairwise Pearson correlations between each sample for the expression levels of all detected mRNAs to determine the reproducibility between our samples. We then used the *heatmap.2* package for graphical representation of correlations, without sample clustering.

#### Gene ontology

Gene ontology was performed using the GO summaries package (V3.11; Kolde and Vilo, 2015) on genes up-regulated (Table S2) and down-regulated in the Olig2-AS (Table S3). Those results were visualized using the REVIGO graphical tool (http://revigo.irb.hr/) and GOplot (V1.0.2; Walter et al., 2015).

#### Tissue collection and processing

Spinal cords at the brachial level were isolated from E13.5 mouse embryos to P7 mouse pups and fixed in 4% paraformaldehyde (PFA, Sigma, CatN#158127) in phosphate-buffered saline (PBS) overnight at 4°C. Tissues were then sectioned at 15 μm to 25 μm using a cryostat (Leica CM1950) after cryoprotection in 20% sucrose (Sigma, CatN# S0389) in PBS and freezing in OCT media (Sakura, CatN: 4583) on the surface of dry ice.

#### In situ RNA hybridization and immunofluorescent staining

Simple or double in situ hybridization and immunofluorescent stainings were performed on transverse sections as previously reported (Ohayon et al., 2019). Digoxigenin- or Fluorescein-labeled antisens RNA probes for *inka2, efnb3*, *fgfr3* and *kcnip3* were synthesized using DIG- or Fluorescein-labelling kit (Roche) according to the manufacturer’s instructions and revealed using either NBT/BCIP (Roche), or Alexa-fluor 488 and 555 Tyramide (Thermo Fisher Scientific). Antibodies used in this study were as follows: rabbit anti-Olig2 (Millipore, CatN#AB9610) and goat anti-Sox10 (Santa Cruz Biotech nology, CatN#sc-17342). Alexa Fluor-488 or Alexa Fluor-647-conjugated secondary antibodies (Thermo Fisher Scientific) were used.

#### Imaging

Confocal images were acquired from tissue sections using Leica SP5 or SP8 confocal microscopes and were always represented as single optical plane sections. Images of simple ISHs were collected with Nikon digital camera DXM1200C on a Nikon Eclipse 80i microscope. Double ISH stainings and double ISH-immunofluorescence stainings were imaged on Leica SP5 and SP8 confocal microscope according to the method of Trinh et al. (Trinh et al., 2007) which allows acquiring in the same optical plane a high resolution confocal image of NBT/BCIP stain in the near infrared range together with the immunofluorescence or TSA signals. Images were processed (size adjustment, luminosity and contrast, and merging or separating layers) using Affinity Photo software (Serif).

### QUANTIFICATION AND STATISTICAL ANALYSIS

#### Figures and representations

*<u>Figure 2F:</u>* Barplot of the relative expression levels (log2FPKM) of CNS cell type specific genes. Graph built using Prism7 (Graphpad)

*<u>Figure S2F:</u>* Barplot of the relative expression levels (log2FPKM) of eGFP and tdTomato transcripts in each replicate. Graph built using Prism7 (Graphpad)

*<u>Figure 3A:</u>* Flowchart representing the treatment and two-step comparative analysis performed on the RNA-seq data. Flowchart built by using the web application draw.io (www.draw.io)

*<u>Figure 3B-E and S3A-C:</u>* Volcano plots were built using the ggplot2 package on the differentially expressed genes representing the log2FC and -log10Padj from the differential analysis DA1, DA2 and DA3.

*<u>Figure 2D-E, 3F-G, S4A-B and S5A-B:</u>* Heatmap representation data (comparisons between the three glial cell populations) were reduced and scaled to generate z-score

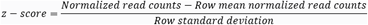

*<u>Figure 5A-B-D:</u>* Treemaps were created using the REVIGO graphical tool (http://revigo.irb.hr/)

*<u>Figure 3C:</u>* Circular visualization of gene-annotation enrichment analysis (GOcircle) with the 245 specifically differentially regulated genes in Olig2-AS (135 genes up-regulated + 110 genes down-regulated).

*<u>Figure 5E-I and 6A-C:</u>* Expression level measurements in Olig2-AS and non-Olig2AS, heatmap are based upon log2FPKM values.

*<u>Figure 5E-I and 6A-C:,</u>* Heatmap representation data (comparison between Olig2-AS and nonOlig2-AS) are based upon log2FC.

All the heatmaps were made with Prism 7 (Graphpad).

#### Table description and cutoffs

Gene lists in supplementary tables were generated as follows:

*<u>Table S2:</u>* genes enriched in Olig2-AS (up1up2), with a log2FC>1 and padj<0.05 both in DA1 and in DA2.

*<u>Table S3:</u>* genes down-regulated in Olig2-AS (dn1dn2), with log2FC<-1 and padj<0.05 both in DA1 and in DA2.

*<u>Table S4:</u>* gene ontology analysis (BP) on genes enriched in Olig2-AS (log2FC>1 and padj<0.05 both in DA1 and in DA2) and in gene depleted in Olig2-AS (log2FC<-1 and padj<0.05 both in DA1 and in DA2).

### Supplemental Items

**Figure S1: Validation of *olig2*-tdTomato transgenic mouse line by immunodetection of Olig2**

**A-C’’’**: Combined detection of Olig2 (blue) and Sox10 (green) on *olig2*-tdTomato transgenic mice at E13.5 (A-A’’’) and P7 (B-C’’’). Vertical sets present successively detection of Olig2, Olig2 and tdT (prime), Sox10 and tdT (double prime) and the merged image (triple prime). Images C-C’’’ show higher magnification of the areas framed in B-B’’’. Asterisks and arrows point to OPC/OL and Olig2-AS, respectively. Scale bars = 100 μm in B-B’’’ and 25 μm in A-A’’’ and C-C’’’.

**Figure S2: Cell isolation strategy and validation of the transcriptome profiling**

**A**: Schematic of the experimental procedure: dissociation and purification of glial cell populations from double transgenic reporter mouse, followed by RNA-seq and analysis. **B**: Principal component analysis (PCA) on all detected mRNAs normalized read count, showing a clear distinction between oligodendrocytes and astroglial subpopulations for mRNA based on the first principal components (PC1). Grouping of the messenger RNA dots suggests separation by cell type. **C**: Heatmap of Pearson correlation between each sample, showing stronger correlation coefficients between samples of the same group than intergroup correlations. **D**: Euclidian unsupervised hierarchical clustering of the standard deviation on normalized expression levels from the 20% most variable mRNAs. **E:**Comparison of eGFP and tdTomato gene expression levels (by log2FPKM values) for each replicate of the three cell populations. All heatmaps are presented as z-score.

**Figure S3: Differential analysis between Olig2-AS, nonOlig2-AS and OPC/OL**

**A-C**: Volcano plots of differentially expressed genes representing the log2FC and -log10Padj from the differential analysis DA1 (A), DA2 (B) and DA3 (C). Red dots represent transcripts with a significant padj<0.05 and a log2FC>0 or a log2FC<0.

**Figure S4: List of genes found up-regulated in Olig2-AS**

**A, B**: Heatmaps depicting the abundance of mRNA corresponding to the 135 genes enriched in Olig2-AS, distinguishing the 39 genes specifically enriched in Olig2-AS (A) and the 96 genes as well enriched in Olig2-AS but also found up-regulated in nonOlig2-AS compared to OPC/OL (DA3). All heatmaps are presented as z-score.

**Figure S5: List of genes found down-regulated in Olig2-AS**

A, B: Heatmaps depicting the abundance of mRNA corresponding to the 110 genes down-regulated in Olig2-AS, distinguishing the 70 genes specifically depleted in the Olig2-AS (A) and the 40 genes as well decreased in Olig2-AS but also found down-regulated in nonOlig2-AS compared to OPC/OL (DA3). All heatmaps are presented as z-score.

**Figure S6: Expression of *inka2* specifically marks astrocyte precursor cells of the developing spinal cord**

A-B’’: Combined detection of Olig2 (green), *inka2* (blue) and*fgfr3* (red) mRNAs in E14.5 embryonic spinal cord. Horizontal sets present successively *fgfr3* and *inka2* staining (A, B), Olig2 and *inka2*staining (A’, B’) and the merged image (A’’, B’’). Images B-B’’ show higher magnification of the areas framed in A-A’’. Asterisks, arrows and arrowheads point to OPC/OL, Olig2-AS and nonOlig2-AS, respectively. Scale bars = 100 μm in A-A’’ and 25 μm in B-B’’.

**Figure S7: Expression pattern of *efnb3***

**A**: Expression profile of *efnb3* mRNA at P7. **B-D’’**: Combined detection of Olig2 (green), and mRNAs encoding for *efnb3* (blue) and *fgfr3* (red). C and D shows higher magnification of the areas framed in B, respectively the grey and white matter. Horizontal sets present successively *fgfr3* and *efnb3* staining (CD), Olig2 and *efnb3* staining (C’-D’) and the merged image (C’’-D’’). Asterisks and arrows point to OPC/OL and Olig2-AS, respectively. Scale bars = 100 μm in A and 50 μm in B and 25 μm in C-C’’ and D-D’’.

**Table S1: FPKM table for the RNA-Seq data**

Expression values for all RNA-Seq data generated in the study, presented as Fragments Per Kilobase of transcript per Million mapped reads (FPKM), for all the glial subpopulations in each replicate.

**Table S2: Significant Olig2-AS enriched genes**

List of genes enriched in Olig2-AS that present a log2FC>1 and padj<0.05 in both DA1 and DA2.

**Table S3: Significant Olig2-AS depleted genes**

List of genes depleted in Olig2-AS that present a log2FC<-1 and padj<0.05 in both DA1 and DA2.

**Table S4: Gene ontology analysis**

Gene ontology analysis (BP) from GOsummaries on genes enriched in Olig2-AS (log2FC>1 and padj<0.05 both in DA1 and in DA2) and in gene depleted in Olig2-AS (log2FC<-1 and padj<0.05 both in DA1 and in DA2). The table indicates the list of GO terms enriched, the list of genes involved in a given GO term, and the p-value. Only terms with p< 0.05 are shown.

## References

Allen, N. J., Bennett, M. L., Foo, L. C., Wang, G. X., Chakraborty, C., Smith, S. J., & Barres, B. A. (2012). Astrocyte glypicans 4 and 6 promote formation of excitatory synapses via GluA1 AMPA receptors. Nature. https://doi.org/10.1038/nature11059

Allen, N. J., & Eroglu, C. (2017). Cell Biology of Astrocyte-Synapse Interactions. In Neuron. https://doi.org/10.1016/j.neuron.2017.09.056

Allen, N. J., & Lyons, D. A. (2018). Glia as architects of central nervous system formation and function. In Science. https://doi.org/10.1126/science.aat0473

Anders, S., Pyl, P. T., & Huber, W. (2014). HTSeq -A Python framework to work with high-throughput sequencing data. BioRxiv. https://doi.org/10.1101/002824

Anderson, C. T., Sheets, P. L., Kiritani, T., & Shepherd, G. M. G. (2010). Sublayer-specific microcircuits of corticospinal and corticostriatal neurons in motor cortex. Nature Neuroscience. https://doi.org/10.1038/nn.2538

Anderson, D., Mehaffey, W. H., Iftinca, M., Rehak, R., Engbers, J. D. T., Hameed, S., Zamponi, G. W., & Turner, R. W. (2010). Regulation of neuronal activity by Cav3-Kv4 channel signaling complexes. Nature Neuroscience. https://doi.org/10.1038/nn.2493

Bansod, S., Kageyama, R., & Ohtsuka, T. (2017). Hes5 regulates the transition timing of neurogenesis and gliogenesis in mammalian neocortical development. Development (Cambridge). https://doi.org/10.1242/dev.147256

Batiuk, M. Y., Martirosyan, A., Wahis, J., de Vin, F., Marneffe, C., Kusserow, C., Koeppen, J., Viana, J. F., Oliveira, J. F., Voet, T., Ponting, C. P., Belgard, T. G., & Holt, M. G. (2020). Identification of region-specific astrocyte subtypes at single cell resolution. Nature Communications. https://doi.org/10.1038/s41467-019-14198-8

Bayraktar, O. A., Fuentealba, L. C., Alvarez-Buylla, A., & Rowitch, D. H. (2015). Astrocyte development and heterogeneity. Cold Spring Harbor Perspectives in Biology, 7(1). https://doi.org/10.1101/cshperspect.a020362

Ben Haim, L., & Rowitch, D. H. (2016). Functional diversity of astrocytes in neural circuit regulation. In Nature Reviews Neuroscience. https://doi.org/10.1038/nrn.2016.159

Boisvert, M. M., Erikson, G. A., Shokhirev, M. N., & Allen, N. J. (2018). The Aging Astrocyte Transcriptome from Multiple Regions of the Mouse Brain. Cell Reports. https://doi.org/10.1016/j.celrep.2017.12.039

Bushong, E. A., Martone, M. E., & Ellisman, M. H. (2004). Maturation of astrocyte morphology and the establishment of astrocyte domains during postnatal hippocampal development. International Journal of Developmental Neuroscience. https://doi.org/10.1016/j.ijdevneu.2003.12.008

Cahoy, J. D., Emery, B., Kaushal, A., Foo, L. C., Zamanian, J. L., Christopherson, K. S., Xing, Y., Lubischer, J. L., Krieg, P. A., Krupenko, S. A., Thompson, W. J., & Barres, B. A. (2008). A transcriptome database for astrocytes, neurons, and oligodendrocytes: A new resource for understanding brain development and function. Journal of Neuroscience. https://doi.org/10.1523/JNEUROSCI.4178-07.2008

Chaboub, L. S., & Deneen, B. (2013). Developmental origins of astrocyte heterogeneity: The final frontier of CNS development. In Developmental Neuroscience. https://doi.org/10.1159/000343723

Chaboub, L. S., Manalo, J. M., Lee, H. K., Glasgow, S. M., Chen, F., Kawasaki, Y., Akiyama, T., Kuo, C. T., Creighton, C. J., Mohila, C. A., & Deneen, B. (2016). Temporal profiling of astrocyte precursors reveals parallel roles for Asef during development and after injury. Journal of Neuroscience, 36(47), 11904–11917. https://doi.org/10.1523/JNEUROSCI.1658-16.2016

Chai, H., Diaz-Castro, B., Shigetomi, E., Monte, E., Octeau, J. C., Yu, X., Cohn, W., Rajendran, P. S., Vondriska, T. M., Whitelegge, J. P., Coppola, G., & Khakh, B. S. (2017). Neural Circuit-Specialized Astrocytes: Transcriptomic, Proteomic, Morphological, and Functional Evidence. Neuron. https://doi.org/10.1016/j.neuron.2017.06.029

Christopherson, K. S., Ullian, E. M., Stokes, C. C. A., Mullowney, C. E., Hell, J. W., Agah, A., Lawler, J., Mosher, D. F., Bornstein, P., & Barres, B. A. (2005). Thrombospondins are astrocyte-secreted proteins that promote CNS synaptogenesis. Cell. https://doi.org/10.1016/j.cell.2004.12.020

Chung, W. S., Allen, N. J., & Eroglu, C. (2015). Astrocytes control synapse formation, function, and elimination. Cold Spring Harbor Perspectives in Biology. https://doi.org/10.1101/cshperspect.a020370

Chung, W. S., Clarke, L. E., Wang, G. X., Stafford, B. K., Sher, A., Chakraborty, C., Joung, J., Foo, L. C., Thompson, A., Chen, C., Smith, S. J., & Barres, B. A. (2013). Astrocytes mediate synapse elimination through MEGF10 and MERTK pathways. Nature. https://doi.org/10.1038/nature12776

Clarke, L. E., Liddelow, S. A., Chakraborty, C., Münch, A. E., Heiman, M., & Barres, B. A. (2018). Normal aging induces A1-like astrocyte reactivity. Proceedings of the National Academy of Sciences of the United States of America. https://doi.org/10.1073/pnas.1800165115

Claus Stolt, C., Rehberg, S., Ader, M., Lommes, P., Riethmacher, D., Schachner, M., Bartsch, U., & Wegner, M. (2002). Terminal differentiation of myelin-forming oligodendrocytes depends on the transcription factor Sox10. Genes and Development. https://doi.org/10.1101/gad.215802

Dessaud, E., McMahon, A. P., & Briscoe, J. (2008). Pattern formation in the vertebrate neural tube: A sonic hedgehog morphogen-regulated transcriptional network. In Development. https://doi.org/10.1242/dev.009324

Dobin, A., Davis, C. a, Schlesinger, F., Drenkow, J., Zaleski, C., Jha, S., Batut, P., Chaisson, M., & Gingeras, T. R. (2012). RNA-STAR: ultrafast universal spliced sequences aligner: Supplementary materials. Bioinformatics (Oxford, England). https://doi.org/10.1093/bioinformatics/bts635

Doyle, J. P., Dougherty, J. D., Heiman, M., Schmidt, E. F., Stevens, T. R., Ma, G., Bupp, S., Shrestha, P., Shah, R. D., Doughty, M. L., Gong, S., Greengard, P., & Heintz, N. (2008). Application of a Translational Profiling Approach for the Comparative Analysis of CNS Cell Types. Cell. https://doi.org/10.1016/j.cell.2008.10.029

Durinck, S., Moreau, Y., Kasprzyk, A., Davis, S., De Moor, B., Brazma, A., & Huber, W. (2005). BioMart and Bioconductor: A powerful link between biological databases and microarray data analysis. Bioinformatics. https://doi.org/10.1093/bioinformatics/bti525

Frank An, W., Bowlby, M. R., Betty, M., Cao, J., Ling, H. P., Mendoza, G., Hinson, J. W., Mattsson, K. I., Strassle, B. W., Trimme, J. S., & Rhodes, K. J. (2000). Modulation of A-type potassium channels by a family of calcium sensors. Nature. https://doi.org/10.1038/35000592

Gokce, O., Stanley, G. M., Treutlein, B., Neff, N. F., Camp, J. G., Malenka, R. C., Rothwell, P. E., Fuccillo, M. V., Südhof, T. C., & Quake, S. R. (2016). Cellular Taxonomy of the Mouse Striatum as Revealed by Single-Cell RNA-Seq. Cell Reports. https://doi.org/10.1016/j.celrep.2016.06.059

Gong, S., Zheng, C., Doughty, M. L., Losos, K., Didkovsky, N., Schambra, U. B., Nowak, N. J., Joyner, A., Leblanc, G., Hatten, M. E., & Heintz, N. (2003). A gene expression atlas of the central nervous system based on bacterial artificial chromosomes. Nature. https://doi.org/10.1038/nature02033

González-Castillo, C., Ortuño-Sahagún, D., Guzmán-Brambila, C., Pallàs, M., & Rojas-Mayorquín, A. E. (2015). Pleiotrophin as a central nervous system neuromodulator, evidences from the hippocampus. Frontiers in Cellular Neuroscience. https://doi.org/10.3389/fncel.2014.00443

Heintz, N. (2004). Gene Expression Nervous System Atlas (GENSAT). In Nature Neuroscience. https://doi.org/10.1038/nn0504-483

Hochstim, C., Deneen, B., Lukaszewicz, A., Zhou, Q., & Anderson, D. J. (2008). Identification of Positionally Distinct Astrocyte Subtypes whose Identities Are Specified by a Homeodomain Code. Cell, 133(3), 510–522. https://doi.org/10.1016/j.cell.2008.02.046

Iwasaki, Y., Yumoto, T., & Sakakibara, S. I. (2015). Expression profiles of inka2 in the murine nervous system. Gene Expression Patterns, 19(1-2), 83–97. https://doi.org/10.1016/j.gep.2015.08.002

Jacobi, E., & von Engelhardt, J. (2018). AMPA receptor complex constituents: Control of receptor assembly, membrane trafficking and subcellular localization. In Molecular and Cellular Neuroscience. https://doi.org/10.1016/j.mcn.2018.05.008

John Lin, C. C., Yu, K., Hatcher, A., Huang, T. W., Lee, H. K., Carlson, J., Weston, M. C., Chen, F., Zhang, Y., Zhu, W., Mohila, C. A., Ahmed, N., Patel, A. J., Arenkiel, B. R., Noebels, J. L., Creighton, C. J., & Deneen, B. (2017). Identification of diverse astrocyte populations and their malignant analogs. Nature Neuroscience. https://doi.org/10.1038/nn.4493

Kelley, K. W., Ben Haim, L., Schirmer, L., Tyzack, G. E., Tolman, M., Miller, J. G., Tsai, H. H., Chang, S. M., Molofsky, A. V., Yang, Y., Patani, R., Lakatos, A., Ullian, E. M., & Rowitch, D. H. (2018). Kir4.1-Dependent Astrocyte-Fast Motor Neuron Interactions Are Required for Peak Strength. Neuron, 98(2), 306–319.e7. https://doi.org/10.1016/j.neuron.2018.03.010

Khakh, B. S., & Sofroniew, M. V. (2015). Diversity of astrocyte functions and phenotypes in neural circuits. Nature Neuroscience. https://doi.org/10.1038/nn.4043

Kiray, H., Lindsay, S. L., Hosseinzadeh, S., & Barnett, S. C. (2016). The multifaceted role of astrocytes in regulating myelination. Experimental Neurology, 283, 541–549. https://doi.org/10.1016/j.expneurol.2016.03.009

Kolde, R., & Vilo, J. (2015). GOsummaries: An R Package for Visual Functional Annotation of Experimental Data. FlOOOResearch. https://doi.org/10.12688/f1000research.6925.1

Kucukdereli, H., Allen, N. J., Lee, A. T., Feng, A., Ozlu, M. I., Conatser, L. M., Chakraborty, C., Workman, G., Weaver, M., Sage, E. H., Barres, B. A., & Eroglu, C. (2011). Control of excitatory CNS synaptogenesis by astrocyte-secreted proteins hevin and SPARC. Proceedings of the National Academy of Sciences of the United States of America. https://doi.org/10.1073/pnas.1104977108

Kuhlbrodt, K., Herbarth, B., Sock, E., Hermans-Borgmeyer, I., & Wegner, M. (1998). Sox10, a novel transcriptional modulator in glial cells. Journal of Neuroscience. https://doi.org/10.1523/jneurosci.18-01-00237.1998

Ledo, F., Link, W. A., Carrión, A. M., Echeverria, V., Mellström, B., & Naranjo, J. R. (2000). The DREAM-DRE interaction: Key nucleotides and dominant negative mutants. In Biochimica et Biophysica Acta - Molecular Cell Research. https://doi.org/10.1016/S0167-4889(00)00092-6

Love, M. I., Anders, S., & Huber, W. (2014). Differential analysis of count data - the DESeq2 package. In Genome Biology. https://doi.org/110.1186/s13059-014-0550-8

Lozzi, B., Huang, T. W., Sardar, D., Huang, A. Y. S., & Deneen, B. (2020). Regionally Distinct Astrocytes Display Unique Transcription Factor Profiles in the Adult Brain. Frontiers in Neuroscience. https://doi.org/10.3389/fnins.2020.00061

Mi, R., Chen, W., & Höke, A. (2007). Pleiotrophin is a neurotrophic factor for spinal motor neurons. Proceedings of the National Academy of Sciences of the United States of America. https://doi.org/10.1073/pnas.0603243104

Miller, S. J., Philips, T., Kim, N., Dastgheyb, R., Chen, Z., Hsieh, Y. C., Daigle, J. G., Datta, M., Chew, J., Vidensky, S., Pham, J. T., Hughes, E. G., Robinson, M. B., Sattler, R., Tomer, R., Suk, J. S., Bergles, D. E., Haughey, N., Pletnikov, M., … Rothstein, J. D. (2019). Molecularly defined cortical astroglia subpopulation modulates neurons via secretion of Norrin. Nature Neuroscience, 22(5), 741–752. https://doi.org/10.1038/s41593-019-0366-7

Mölders, A., Koch, A., Menke, R., & Klöcker, N. (2018). Heterogeneity of the astrocytic AMPA-receptor transcriptome. GLIA. https://doi.org/10.1002/glia.23514

Molofsky, A. V., Kelley, K. W., Tsai, H. H., Redmond, S. A., Chang, S. M., Madireddy, L., Chan, J. R., Baranzini, S. E., Ullian, E. M., & Rowitch, D. H. (2014). Astrocyte-encoded positional cues maintain sensorimotor circuit integrity. Nature, 509(7499), 189–194. https://doi.org/10.1038/nature13161

Morel, L., Chiang, M. S. R., Higashimori, H., Shoneye, T., Iyer, L. K., Yelick, J., Tai, A., & Yang, Y. (2017). Molecular and functional properties of regional astrocytes in the adult brain. Journal of Neuroscience. https://doi.org/10.1523/JNEUROSCI.3956-16.2017

Muroyama, Y., Fujiwara, Y., Orkin, S. H., & Rowitch, D. H. (2005). Specification of astrocytes by bHLH protein SCL in a restricted region of the neural tube. Nature. https://doi.org/10.1038/nature04139

Ohayon, D., Escalas, N., Cochard, P., Glise, B., Danesin, C., & Soula, C. (2019). Sulfatase 2 promotes generation of a spinal cord astrocyte subtype that stands out through the expression of Olig2. GLIA, 67(8). https://doi.org/10.1002/glia.23621

Pestana, F., Edwards-Faret, G., Belgard, T. G., Martirosyan, A., & Holt, M. G. (2020). No longer underappreciated: The emerging concept of astrocyte heterogeneity in neuroscience. In Brain Sciences. https://doi.org/10.3390/brainsci10030168

Pfrieger, F. W., & Ungerer, N. (2011). Cholesterol metabolism in neurons and astrocytes. Progress in Lipid Research, 50(4), 357–371. https://doi.org/10.1016/j.plipres.2011.06.002

Pringle, N. P., Yu, W. P., Howell, M., Colvin, J. S., Ornitz, D. M., & Richardson, W. D. (2003). Fgfr3 expression by astrocytes and their precursors: Evidence that astrocytes and oligodendrocytes originate in distinct neuroepithelial domains. In Development. https://doi.org/10.1242/dev.00184

Saunders, A., Macosko, E. Z., Wysoker, A., Goldman, M., Krienen, F. M., de Rivera, H., Bien, E., Baum, M., Bortolin, L., Wang, S., Goeva, A., Nemesh, J., Kamitaki, N., Brumbaugh, S., Kulp, D., & McCarroll, S. A. (2018). Molecular Diversity and Specializations among the Cells of the Adult Mouse Brain. Cell. https://doi.org/10.1016/j.cell.2018.07.028

Simard, M., & Nedergaard, M. (2004). The neurobiology of glia in the context of water and ion homeostasis. Neuroscience. https://doi.org/10.1016/j.neuroscience.2004.09.053

Sofroniew, M. V., & Vinters, H. V. (2010). Astrocytes: Biology and pathology. In Acta Neuropathologica. https://doi.org/10.1007/s00401-009-0619-8

Stogsdill, J. A., Ramirez, J., Liu, D., Kim, Y. H., Baldwin, K. T., Enustun, E., Ejikeme, T., Ji, R. R., & Eroglu, C. (2017). Astrocytic neuroligins control astrocyte morphogenesis and synaptogenesis. Nature, 551(7679), 192–197. https://doi.org/10.1038/nature24638

Sun, W., McConnell, E., Pare, J. F., Xu, Q., Chen, M., Peng, W., Lovatt, D., Han, X., Smith, Y., & Nedergaard, M. (2013). Glutamate-dependent neuroglial calcium signaling differs between young and adult brain. Science. https://doi.org/10.1126/science.1226740

Tekkök, S. B., Brown, A. M., Westenbroek, R., Pellerin, L., & Ransom, B. R. (2005). Transfer of glycogen-derived lactate from astrocytes to axons via specific monocarboxylate transporters supports mouse optic nerve activity. Journal of Neuroscience Research. https://doi.org/10.1002/jnr.20573

Trinh, L. A., McCutchen, M. D., Bonner-Fraser, M., Fraser, S. E., Bumm, L. A., & McCauley, D. W. (2007). Fluorescent in situ hybridization employing the conventional NBT/BCIP chromogenic stain. BioTechniques. https://doi.org/10.2144/000112476

Uchigashima, M., Cheung, A., Suh, J., Watanabe, M., & Futai, K. (2019). Differential expression of neurexin genes in the mouse brain. Journal of Comparative Neurology, 527(12), 1940–1965. https://doi.org/10.1002/cne.24664

Walter, W., Sánchez-Cabo, F., & Ricote, M. (2015). GOplot: An R package for visually combining expression data with functional analysis. Bioinformatics. https://doi.org/10.1093/bioinformatics/btv300

Zeisel, A., Hochgerner, H., Lönnerberg, P., Johnsson, A., Memic, F., van der Zwan, J., Häring, M., Braun, E., Borm, L. E., La Manno, G., Codeluppi, S., Furlan, A., Lee, K., Skene, N., Harris, K. D., Hjerling-Leffler, J., Arenas, E., Ernfors, P., Marklund, U., & Linnarsson, S. (2018). Molecular Architecture of the Mouse Nervous System. Cell. https://doi.org/10.1016/j.cell.2018.06.021

Zeisel, A., Moz-Manchado, A. B., Codeluppi, S., Lönnerberg, P., Manno, G. La, Juréus, A., Marques, S., Munguba, H., He, L., Betsholtz, C., Rolny, C., Castelo-Branco, G., Hjerling-Leffler, J., & Linnarsson, S. (2015). Cell types in the mouse cortex and hippocampus revealed by single-cell RNA-seq. Science. https://doi.org/10.1126/science.aaa1934

Zhang, Y., & Barres, B. A. (2010). Astrocyte heterogeneity: An underappreciated topic in neurobiology. In Current Opinion in Neurobiology. https://doi.org/10.1016/j.conb.2010.06.005

Zhang, Y., Chen, K., Sloan, S. A., Bennett, M. L., Scholze, A. R., O’Keeffe, S., Phatnani, H. P., Guarnieri, P., Caneda, C., Ruderisch, N., Deng, S., Liddelow, S. A., Zhang, C., Daneman, R., Maniatis, T., Barres, B. A., & Wu, J. Q. (2014). An RNA-sequencing transcriptome and splicing database of glia, neurons, and vascular cells of the cerebral cortex. Journal of Neuroscience. https://doi.org/10.1523/JNEUROSCI.1860-14.2014

Zhou, Q., Wang, S., & Anderson, D. J. (2000). Identification of a novel family of oligodendrocyte lineage-specific basic helix-loop-helix transcription factors. Neuron. https://doi.org/10.1016/S0896-6273(00)80898-3

