## Supplementary figures and images for "Transcriptome profiling of the Olig2-expressing astrocyte subtype reveals their unique molecular signature"

### Supplemental Figure 1

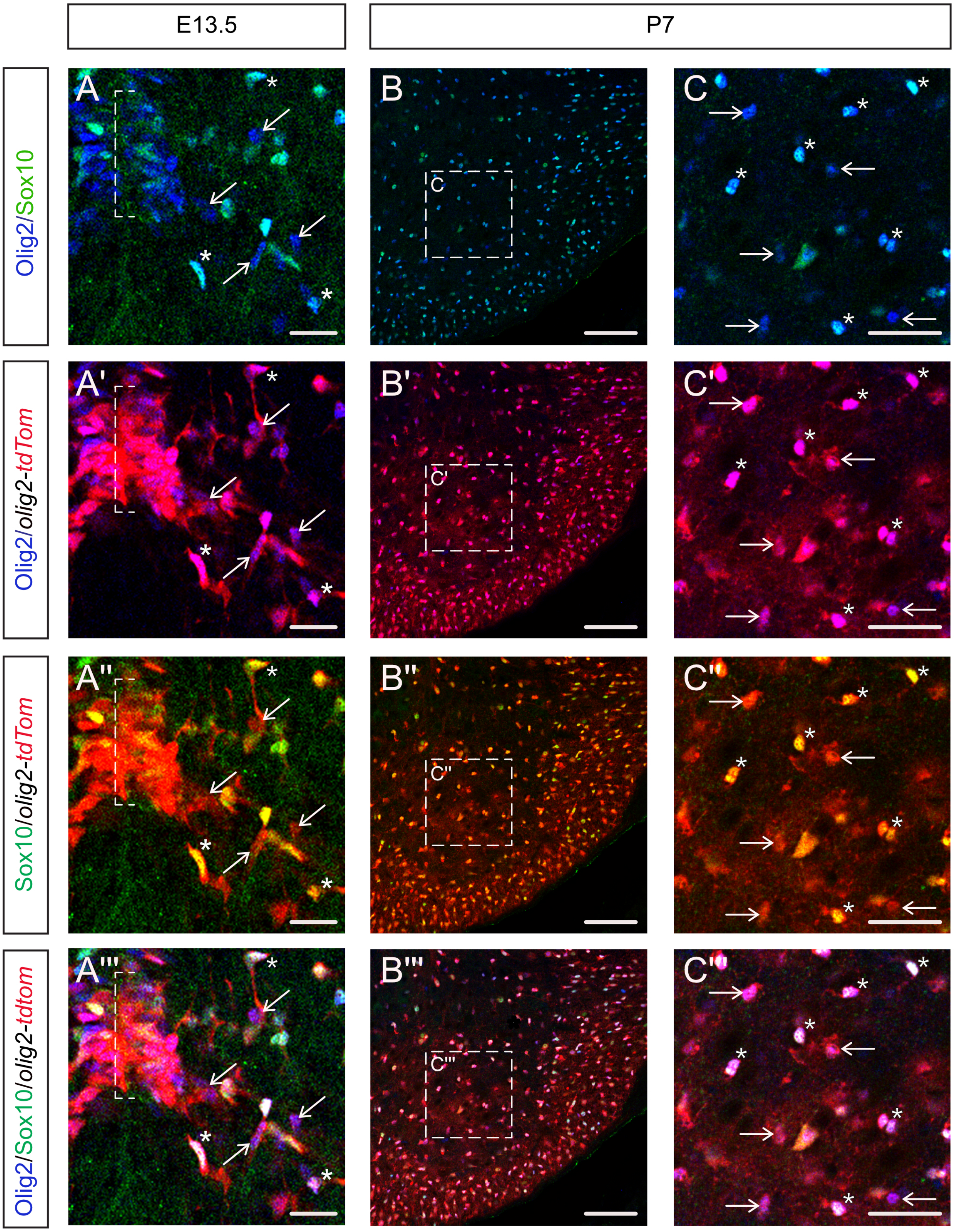

### Supplemental Figure 2

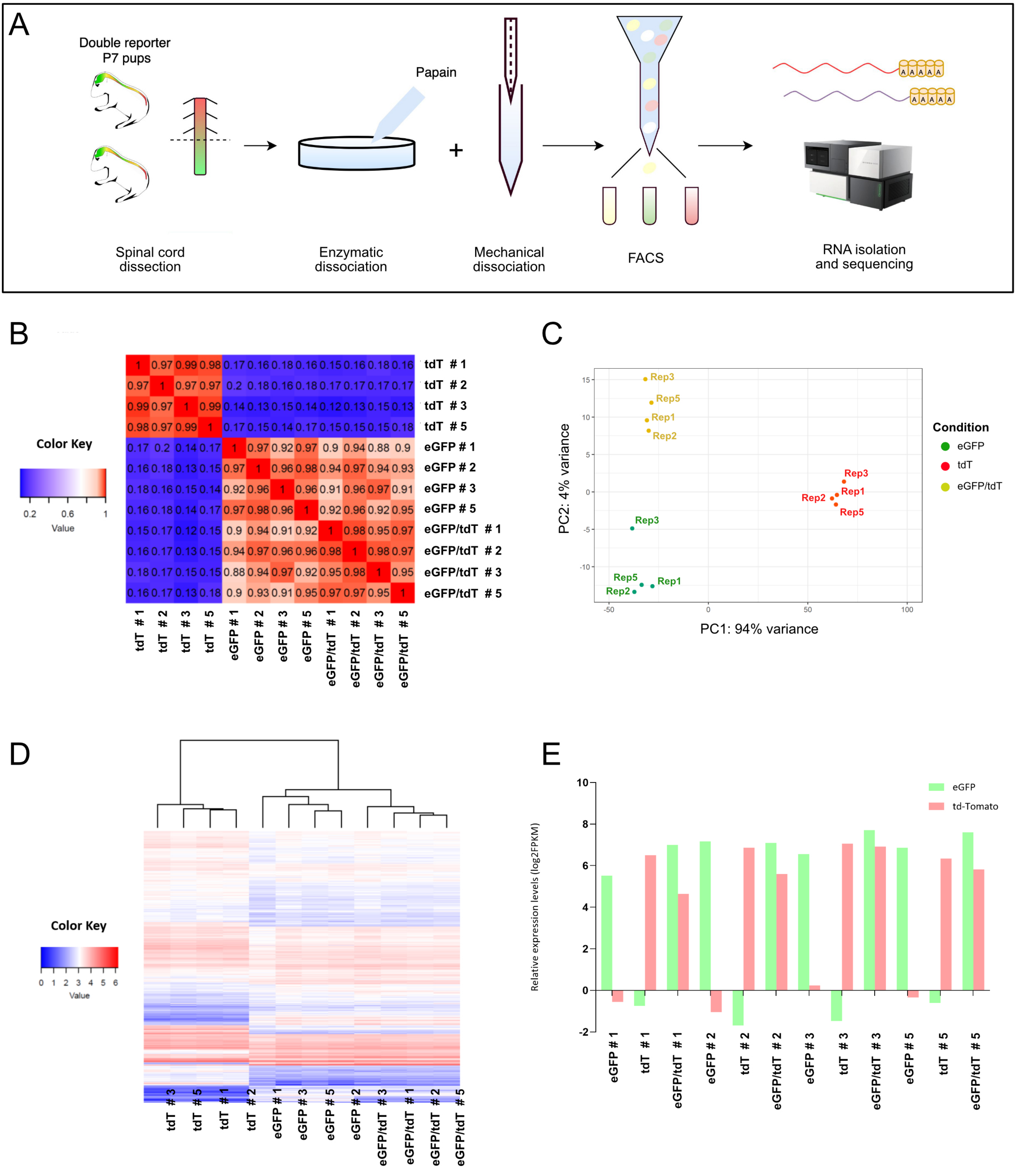

### Supplemental Figure 3

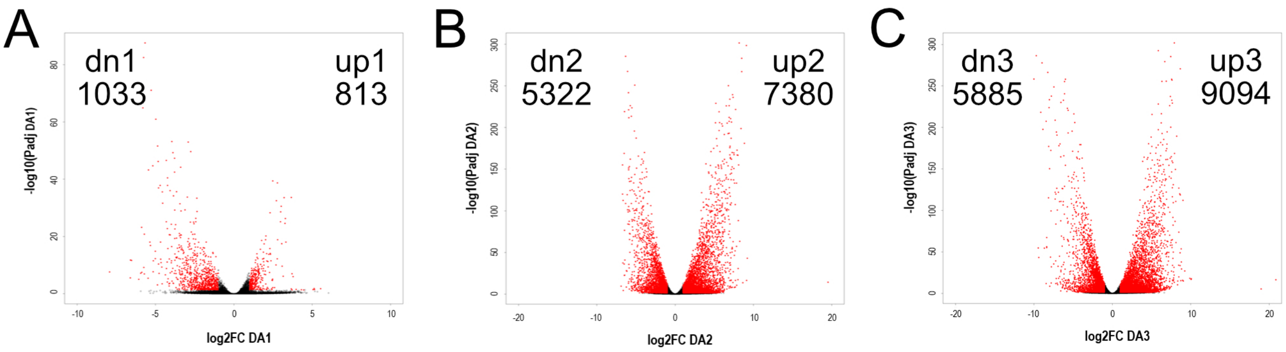

### Supplemental Figure 4

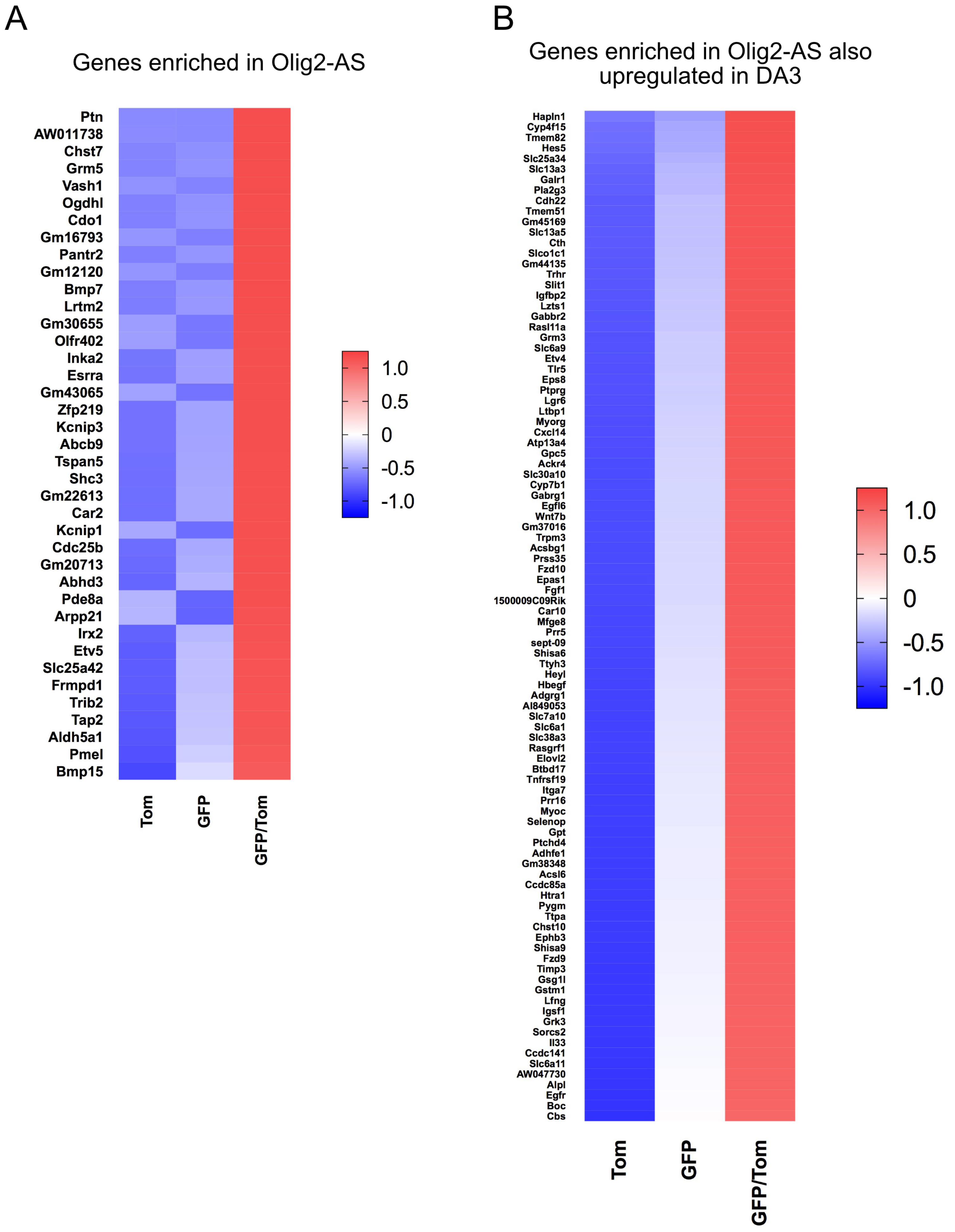

### Supplemental Figure 5

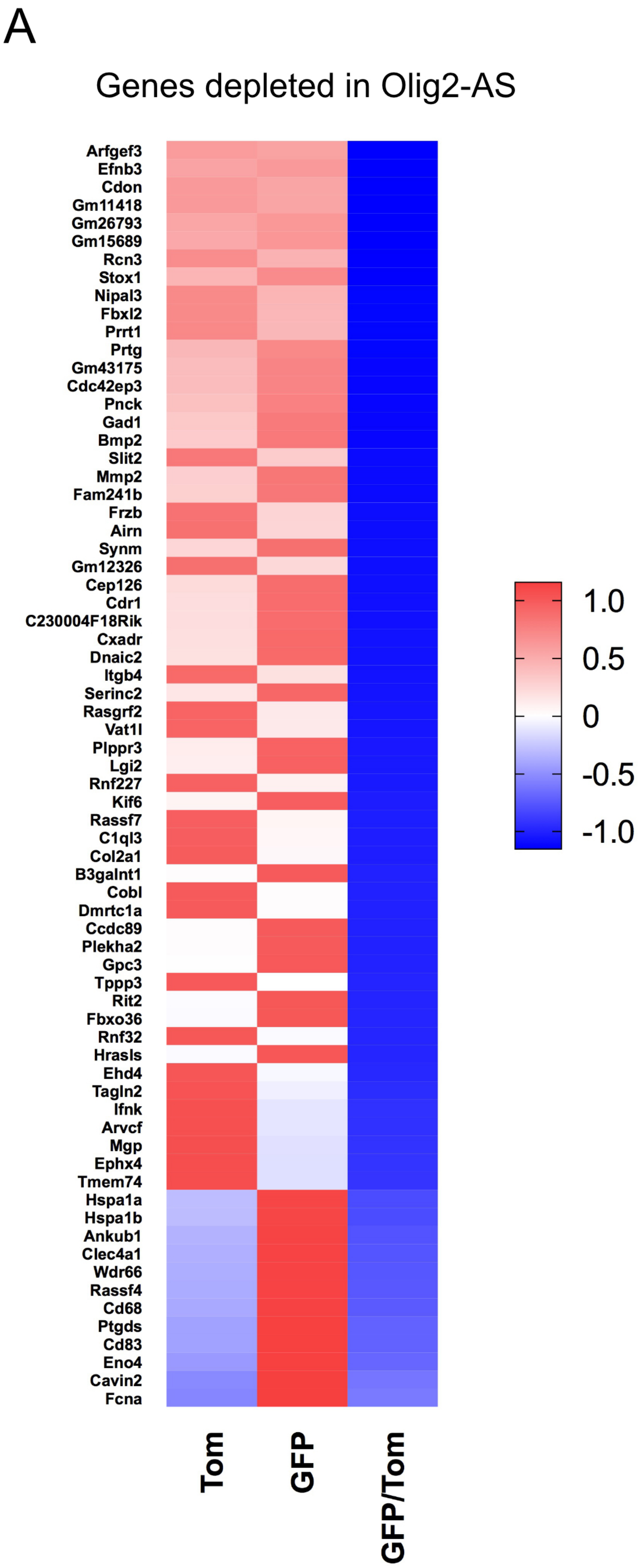

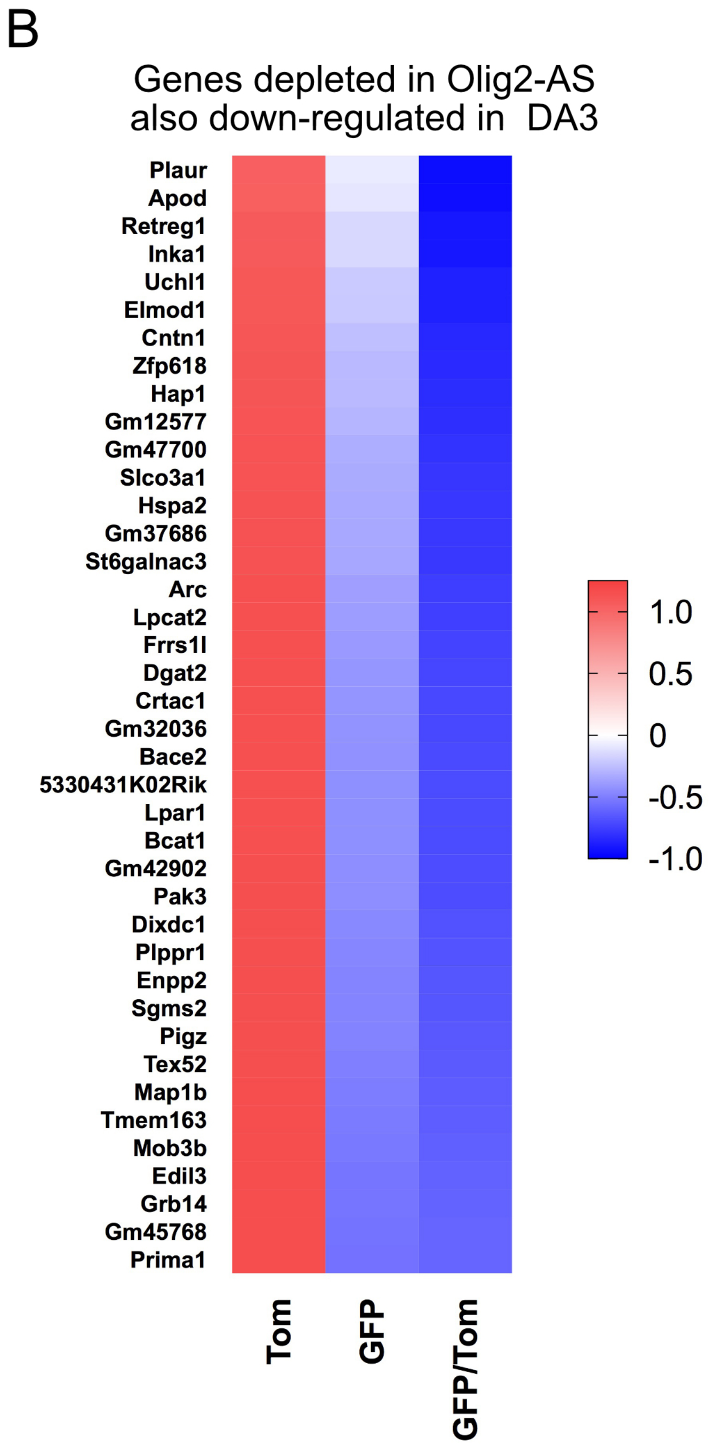

### Supplemental Figure 6

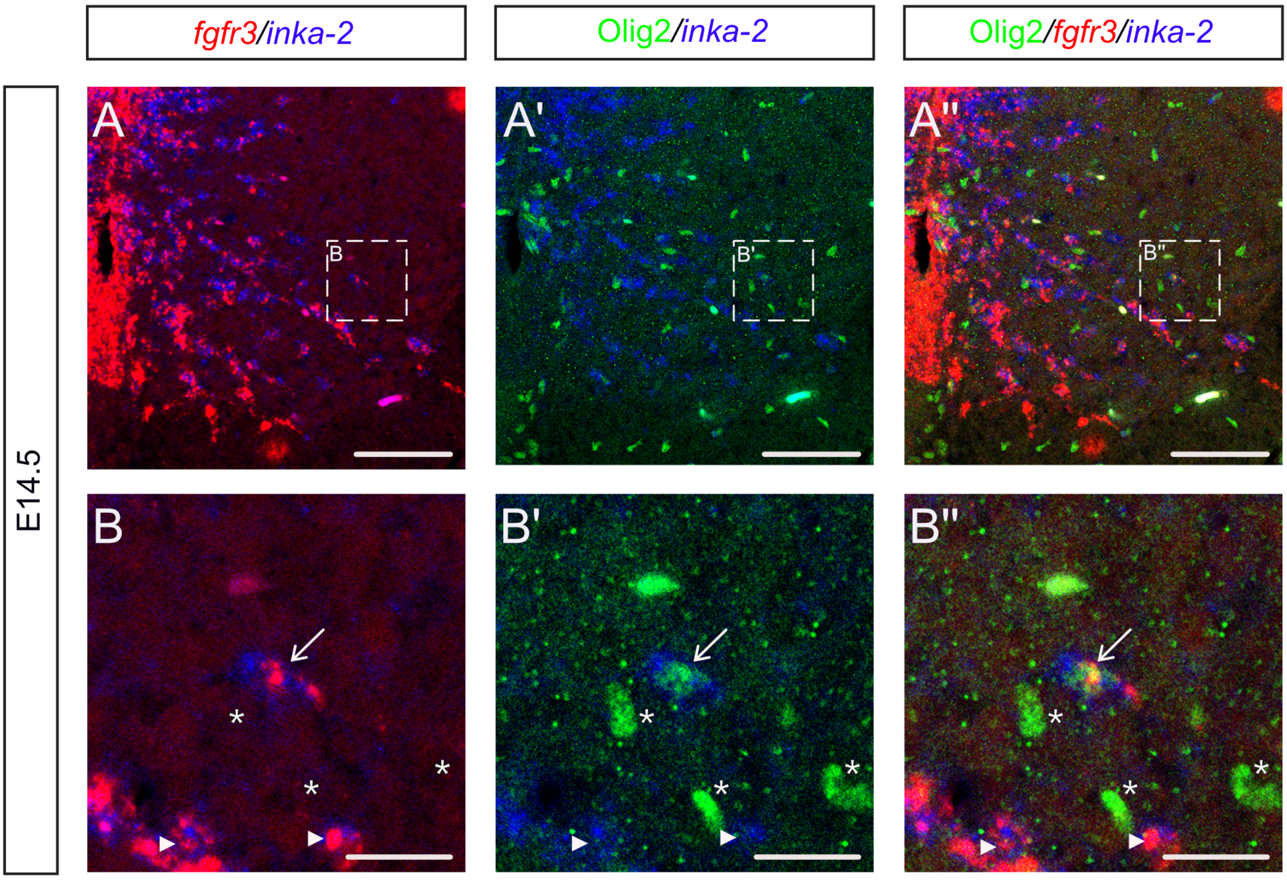


Figure S6

### Supplemental Figure 7

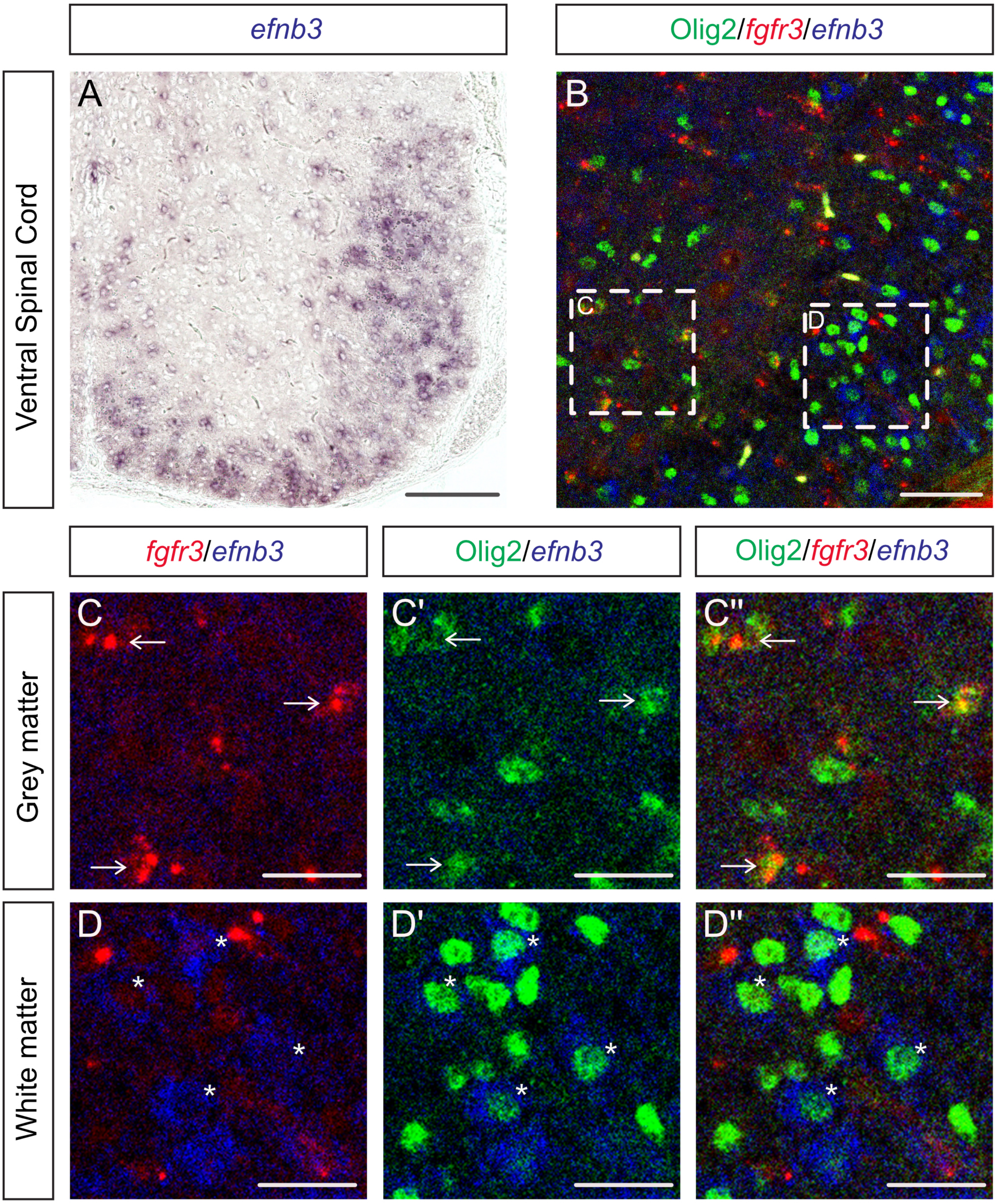
